# A feed-forward loop between niche adenosine and Gzmk⁺ CD8 T cells propagates systemic inflammaging

**DOI:** 10.64898/2026.03.18.712515

**Authors:** Lin Guo, Rui Zheng, Qianqian Zhan, Xin Yu, Qi Zhang, Jiaxi Yao, Xiaoqing Tan, Xiaoting Li, Ou Zhang, Peiyun Liu, Chenghan Wang, Ye Yao, Rui Ma, Xiao Wu, Chenyuan Wang, Haonan Zhou, Yijun Sun, Kai Xiong, Linlin Li, Huajun Xu, Mingzhu Zheng, Jun Jin, Yan Wu, Tao Liu, Zhuo Wang, Yang Liu, Zhan-you Wang, Wenqiang Cao

## Abstract

The causal link between the aging microenvironment and T cell aging remains elusive. Here, we demonstrate that adenosine within aging tissues actively reprograms CD8+ T cells into a pro-aging Granzyme K^+^ (Gzmk+) population. Mechanistically, senescent cells create an adenosine-rich niche via p16-dependent CD39 upregulation, triggering A2aR signaling to induce Gzmk^+^ T cell differentiation. Once released, Gzmk promotes systemic inflammaging through PAR1 and complement activation. Crucially, targeting this axis—either via genetic Gzmk ablation or pharmacological A2aR blockade—reverses multi-organ aging phenotypes and significantly extends healthy lifespan in mice. Human analysis reveals age-dependent Gzmk^+^ T cell accumulation in multi organs, while coffee intake (an A2aR antagonist) inversely correlates with plasma Gzmk levels. Our findings uncover how metabolic niche changes drive T cell aging and establish the adenosine-Gzmk axis as a pivotal therapeutic target for combating age-related diseases.

## Introduction

Organismal aging is characterized by a progressive erosion of tissue homeostasis and function, closely linked to chronic, sterile, low-grade inflammation-- termed “inflammaging”(*1–4*). A pivotal driver of this process is the accumulation of senescent cells. Despite losing proliferative capacity, these cells remain metabolically active and secrete a pathogenic cocktail of cytokines, chemokines, and proteases known as the senescence-associated secretory phenotype (SASP)(*5–8*). While the accumulation of senescent cells is widely recognized as a primary hallmark of aging, the upstream regulatory networks that orchestrate their accumulation and the persistence and propagate this inflammatory milieu remain incompletely understood(*9*).

The immune system undergoes profound remodeling during aging, collectively referred as immunosenescence(*10*). Emerging evidence indicates that aged immune cells are not merely victims of systemic aging but active drivers of multiorgan dysfunction(*11, 12*). Among these key features, T cell aging represents a critical node; while age-associated T cell dysfunction classically increases vulnerability to infections and malignancies(*13–15*). A recent study demonstrates that selective depletion of *Tfam* in T cells accelerates multiple-organ aging(*16*), highlighting a a causal role for T cells in actively dictating organismal lifespan.

Canonically, T cells maintain tissue health by surveilling and clearing senescent cells(*17–22*), As T cell function wanes with age, this immune surveillance collapses, facilitating the accumulation of senescent cells and acceleraing aging and age-related pathologies(*18, 21–23*).Furthermore, aged immune cells can actively induce paracrine senescence(*11*). However, the specific T cell subsets that dominate this pathogenic paracrine signaling, and the precise molecular mechanisms by which the aging tissue microenvironment corrupts T cells to create a causative niche for systemic aging, remain central unresolved questions.

Recent single-cell transcriptomic analyses have revealed a highly conserved, age-associated CD8 T cells (Taa cells) population characterized by the high expression of Gzmk. These cells expand in aged mice tissues and peripheral blood mononuclear cells (PBMCs) from older humans(*24*). Unlike their Granzyme B (Gzmb)-expressing counterparts, which execute targeted cytotoxicity, Gzmk^+^ CD8 T cells have been increasingly implicated in fueling chronic inflammation(*25–30*). Although they are recognized as conserved a hallmark of inflammaging (*24, 31, 32*), their precise physiological role in organismal aging remains ambiguous. In parallel, the aging microenvironment undergoes metabolic rewiring, characterized by the accumulation of extracellular metabolites. Whether these metabolic cues instruct the differentiation of Gzmk^+^ T cells, and whether extracellular Gzmk exerts non-cytotoxic functions to actively propagate tissue aging has yet to be determined.

In this study, we decipher the ontogeny and systemic pathogenicity of age-associated Gzmk^+^ CD8 T cells. We demonstrate that the aging tissue microenvironment actively orchestrates the generation of these cells through a novel metabolic-immune axis: senescence-associated upregulation of CD39 drives the local accumulation of extracellular adenosine, which acts via the A2aR receptor on CD8⁺ T cells to induce the master transcription factor Eomes. This adenosine–Eomes axis, synergizing with PD-1/PD-L1 signaling, locks T cells into a stable Gzmk⁺ state. Once secreted, Gzmk functions not as a cytotoxic killer, but as a potent pro-senescence protease. By cleaving the extracellular receptor PAR1 and complement component C3, Gzmk triggers a vicious feed-forward loop of cellular senescence and systemic inflammaging. Highlighting the robust translational potential of this axis, we validate the accumulation of GZMK⁺ CD8⁺ T cells in normal aged human tissues and report a striking inverse correlation between habitual caffeine (a natural A2aR antagonist) consumption and circulating GZMK levels in a human cohort. Crucially, targeted disruption of this axis—either via genetic *Gzmk* ablation or pharmacological inhibition of A2aR or Gzmk—reverses age-related phenotypes and rejuvenates tissue function. Together, our findings delineate a unifying pathogenic metabolic-immune axis, providing a new conceptual framework and actionable therapeutic targets to combat systemic aging.

## Results

### Gzmk^+^ CD8⁺ T cells are generated locally in response to organ-specific aging

While the aged immune system exhibits both systemic and organ-specific remodeling, the precise developmental origins of aging-associated Gzmk⁺ CD8 T cells remain incompletely understood. A previous murine study has proposed that aged adipose tissue serves as a primary reservoir driving the generation and systemic expansion of these cells during(*32*). However, the robust accumulation of GZMK⁺ CD8⁺ T cells has also been documented in several human inflammatory diseases (*27–30*), hinting at an adipose-independent, tissue-autonomous induction mechanism. To determine whether GZMK⁺ CD8 T cells accumulate in normal human tissues during natural aging, we profiled tonsil, kidney, and skin specimens from young and older individuals. Flow cytometric and multicolored immunohistochemical analyses revealed a significant, age-dependent enrichment of GZMK⁺ CD8 T cells across diverse immune and non-immune human tissues, accompanied by elevated global GZMK protein levels (Fig. 1A-F). This pattern closely mirrored our observations in aging murine tissues (fig. S1A). Thus, the progressive accumulation of GZMK⁺ CD8 T cells is a conserved hallmark of natural aging in human peripheral tissues.

**Figure 1.**
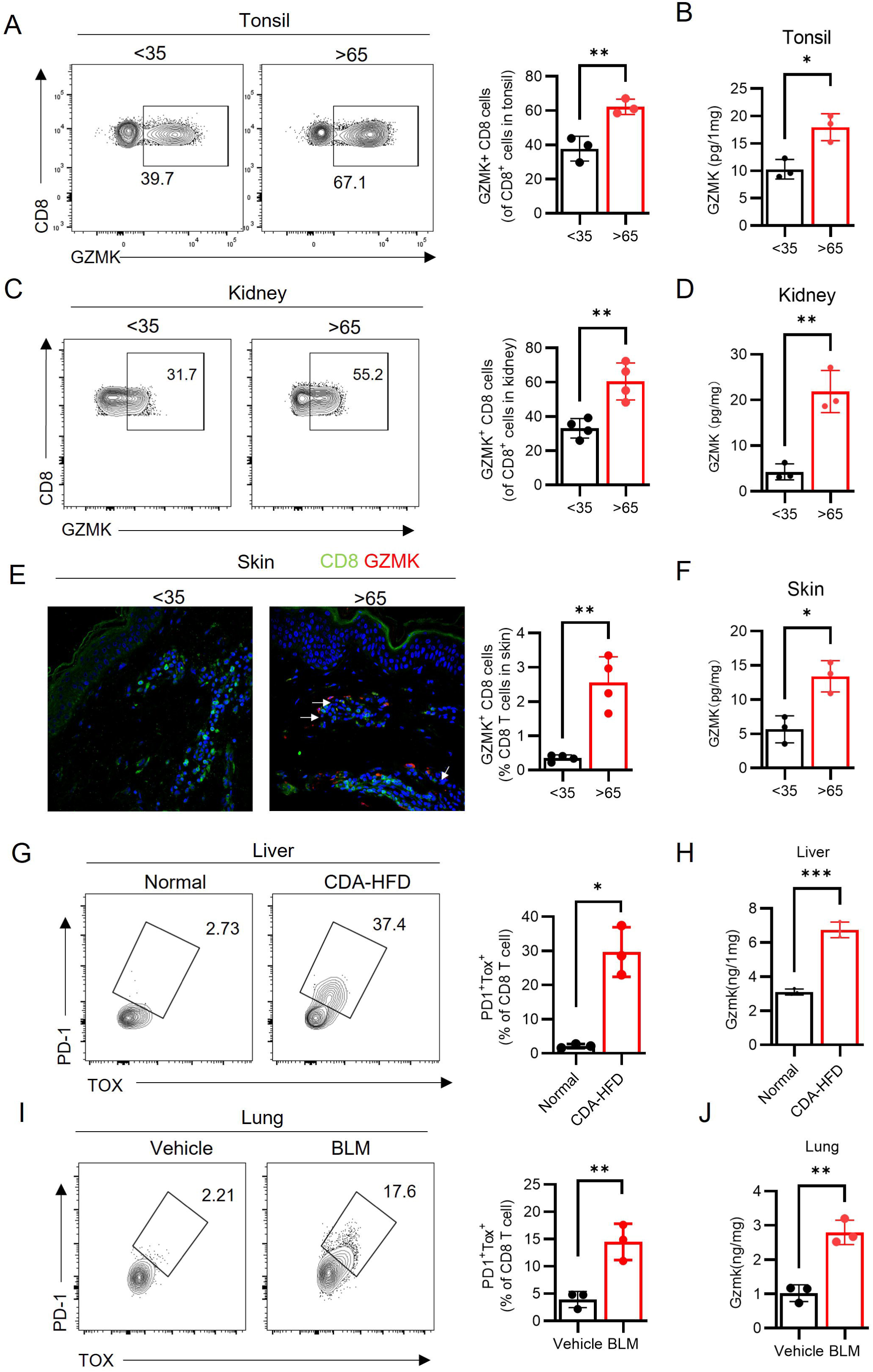
Gzmk^+^ CD8⁺ T cells are generated locally in response to organ-specific aging. (A, C) Analysis of CD8 T cells in tonsil (A), kidney (C) from young (<35 years old) or old (>65 years old) individuals by flow cytometry; (G) multi-color immunohistology staining of GZMK and CD8 in skins from young and old individuals. Flow cytometry analysis of CD8 T cells the liver (G) from control or NASH mice; the lung (I) from control or BLM-treated mice. (B, D, F, H, J) Gzmk levels in indicated tissues were analyzed by ELISA.

The pronounced presence of GZMK⁺ CD8⁺ T cells across disparate aged and inflamed tissues led us to hypothesize that these cells are induced locally by the aging tissue microenvironment rather than exclusively recruited from systemic pools. To test this, we employed two independent tissue-specific aging models. First, we utilized nonalcoholic steatohepatitis (NASH) in mice by feeding a choline-deficient, L-amino-acid-defined, high-fat diet (CDA-HFD)(*22, 33*), a pathology intimately linked to hepatic cellular senescence(*34–36*). This localized senescent environment drove a >10-fold increase in the frequency of Taa cells strictly within the liver, while splenic frequencies remained unperturbed (Fig. 1G-H and fig. S2B). Second, we induced lung-specific cellular senescence via airway administration of bleomycin (BLM)(*18, 37*). Consistent with the NASH model, Taa cells expanded markedly within the senescent lung microenvironment, whereas their baseline frequency in the spleen remained unaltered (Fig.1I and fig. S2C). Collectively, these targeted *in vivo* models demonstrate that senescent tissue niches possess the intrinsic capacity to locally instruct the generation and expansion of pathogenic Gzmk⁺ CD8 T cells.

### Adenosine accumulation in aged tissues drives Gzmk⁺ CD8 T cell differentiation via A2aR–Eomes signaling

To identify conserved tissue-derived metabolites responsible for the induction of Gzmk⁺ CD8 T cells across disparate organs, we performed matrix-assisted laser desorption/ionization mass spectrometry imaging (MALDI-MSI; hereafter referred to as IMS). Spatially resolved mass spectra were acquired from frozen liver sections of young, aged, and NASH-model mice, alongside kidney specimens from young and elderly human donors. This untargeted metabolomic profiling identified 112 metabolites that were commonly elevated in the livers of aged and NASH mice, as well as in elderly human kidneys (fig. S2A). Among these metabolites, adenosine—identified by m/z 268.104 ± 0.05— emerged as a prime candidate, given that its metabolites are associated with inflammaging(*38, 39*), and its capacity to induce the co-inhibitory receptor expression on CD8 T cells(*40, 41*), a core phenotypic feature of Gzmk^+^ CD8 T cells. Further analysis confirmed significantly higher adenosine peaks within these aged and pathological tissues (fig. S2B). Visualization of IMS data demonstrated broadly distributed adenosine signatures throughout the full tissue architecture of both aged mouse livers and human kidneys (Fig. 2A and fig. S3A). These localized spatial increases were corroborated by ELISA, which demonstrated a marked, age-dependent accumulation of adenosine across murine liver, lung, and adipose tissues, as well as in human tonsil, skin, and kidney samples (Fig. 2B, C). Thus, elevated tissue adenosine represents a conserved biochemical hallmark of organ aging.

**Figure 2.**
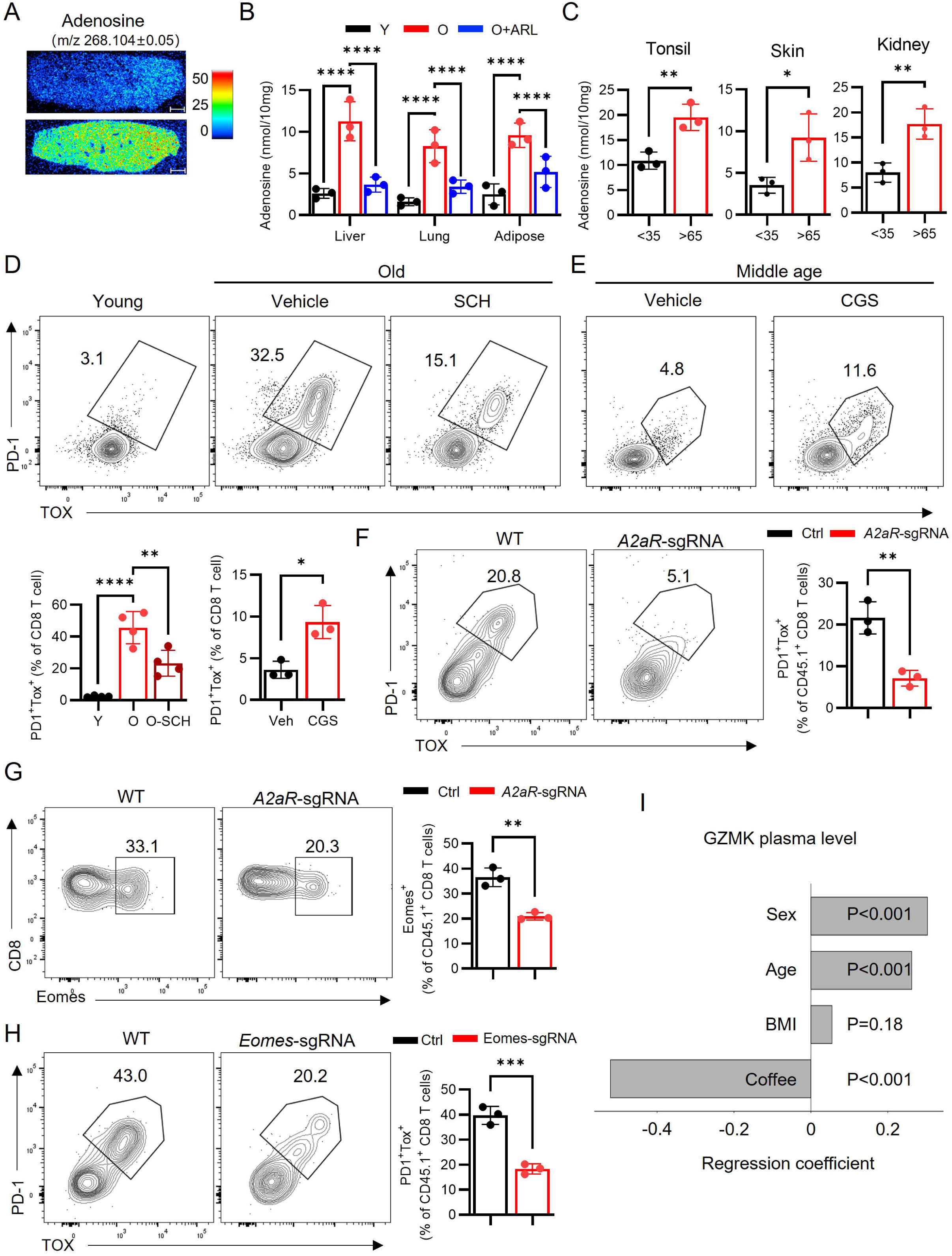
Adenosine accumulation in aged tissues drives Gzmk⁺ CD8 T cell differentiation via A2aR–Eomes signaling. (A) Signaling of adenosine by IMS in liver slides from young or old mice. (B) Adenosine was measured in liver, lung and adipose from young, old or ARL-treated old mice (for one month). (C) Adenosine level in indicated tissues from young and old individuals. PD1^+^ TOX^+^ (Taa) cells in spleen from young, old or SCH-treated old mice (D), from vehicle or CGS-treated middle-aged mice (12months) (E) analyzed by flow cytometry. CD8 T cells from CD45.1 Cas9-expressing mice were electroporated with control or *A2aR* sgRNA and were adoptively transferred into old mice. After one month, PD1^+^ TOX^+^ cells (F), expression of Eomes (G) in CD45.1 CD8 T cells in spleen were analyzed by flow cytometry. (H) Experiments were performed same with Figure 2F using control or *Eomes* sgRNA, PD1^+^ TOX^+^ cells among CD45.1^+^ donor cells were analyzed by flow cytometry. (I) Multiple regression analysis was conducted on GZMK plasma levels (*n* = 518) and coffee intake or not (adjusted for age, sex and BMI).

Adenosine exerts its immunomodulatory effects primarily through the A2aR, the predominant adenosine receptor subtype expressed on T cells(*42–44*). To determine whether this adenosine-rich microenvironment actively drives Gzmk⁺ CD8⁺ T cell generation, we treated aged mice with the selective A2aR inhibitor SCH□442416. Pharmacological blockade of A2aR over one month profoundly reduced both the frequency of Gzmk⁺ CD8 T cells in the spleen and liver (Fig. 2D and fig. S4A). Conversely, *in vivo* administration of the A2aR agonist CGS21680 was sufficient to expand the Gzmk⁺ CD8 T cell pool in these tissues in middle-aged mice (Fig. 2E and fig. S4B). To definitively establish the T cell-intrinsic requirement of A2aR signaling, we adoptively transferred *Adora2a*-deleted (sgRNA-electroporated) CD8⁺ T cells from CD45.1⁺ Cas9-expressing mice into aged wild-type recipients. Genetic ablation of A2aR markedly decreased the generation of Gzmk⁺ CD8⁺ T cells within the aged host microenvironment (Fig. 2F and fig. S5A, B). These data demonstrate that T cell-intrinsic A2aR signaling is required to translate age-associated adenosine accumulation into pathogenic Gzmk⁺ CD8 T cell differentiation.

Previous transcriptomic profiling has linked the Gzmk⁺ CD8⁺ T cell state to elevated expression of the transcription factor Eomes, which has been shown to directly transactivate the *Gzmk* locus in human CD4⁺ T cells (*24, 45, 46*). We therefore hypothesized that Eomes operates downstream of A2aR as the master transcriptional regulator of Gzmk^+^ CD8 T cells. Indeed, A2aR-deficiency in CD8⁺ T cells significantly decreased Eomes expression following adoptive transfer into aged hosts (Fig. 2G). whereas CGS21680 treatment robustly induced Eomes expression in middle-aged mice (fig. S5C). Direct CRISPR/Cas9-mediated deletion of *Eomes* in donor CD8⁺ T cells prior to adoptive transfer into aged recipients severely impaired their subsequent differentiation into Gzmk⁺ CD8 T cells (Fig. 2H and fig. S5D, E), establishing Eomes as the indispensable transcriptional link between microenvironmental adenosine and the differentiation of Gzmk+ CD8 T cells.

To evaluate the translational relevance of this adenosine–A2aR–GZMK axis in human aging, we capitalized on the fact that caffeine is a natural structural analog of adenosine and a widely consumed A2aR antagonist. In a human cohort comprising 518 individuals, we observed a progressive, age-dependent escalation in plasma GZMK levels, consistent with the systemic accumulation of GZMK⁺ CD8 T cells (Fig. 2I). Strikingly, multiple regression analysis—adjusted for age, sex, and body mass index (BMI)—revealed a robust inverse correlation between regular coffee intake and circulating GZMK concentrations (Fig. 2I). These epidemiological findings compellingly reinforce our murine mechanistic data, supporting a model wherein blockade of senescence-associated adenosine signaling attenuates the generation of pathogenic GZMK⁺ CD8⁺ T cells in humans.

### p16-dependent CD39 induction in senescent cells drives adenosine accumulation and promotes Gzmk⁺ CD8 T cell differentiation in aged tissues

As shown above, adenosine is required for the differentiation of Gzmk⁺ CD8 T cells in aged tissues; thus, manipulating adenosine production may represent an effective strategy to reduce the accumulation of these cells. Extracellular adenosine is generated primarily from ATP through sequential enzymatic conversion by CD39 and CD73(*47*). IMS analysis revealed a mild increase in ATP levels but a marked increase in ADP/AMP and adenosine in aged liver (fig. S3B), indicative of elevated CD39 and CD73 enzymatic activity in aged tissue. We next examined the expression profiles of CD39 and CD73 during aging. Expression of CD39 on CD45⁻ non-immune cells was significantly higher in aged tissues, whereas CD73 levels remained largely unchanged (Fig. 3A and fig. S6A). These findings were confirmed by histological staining in livers from young or old mice (Fig. 3B). Increased CD39 expression on CD45⁻ cells was also observed in skin samples from older human donors (Fig. 3C).

**Figure 3.**
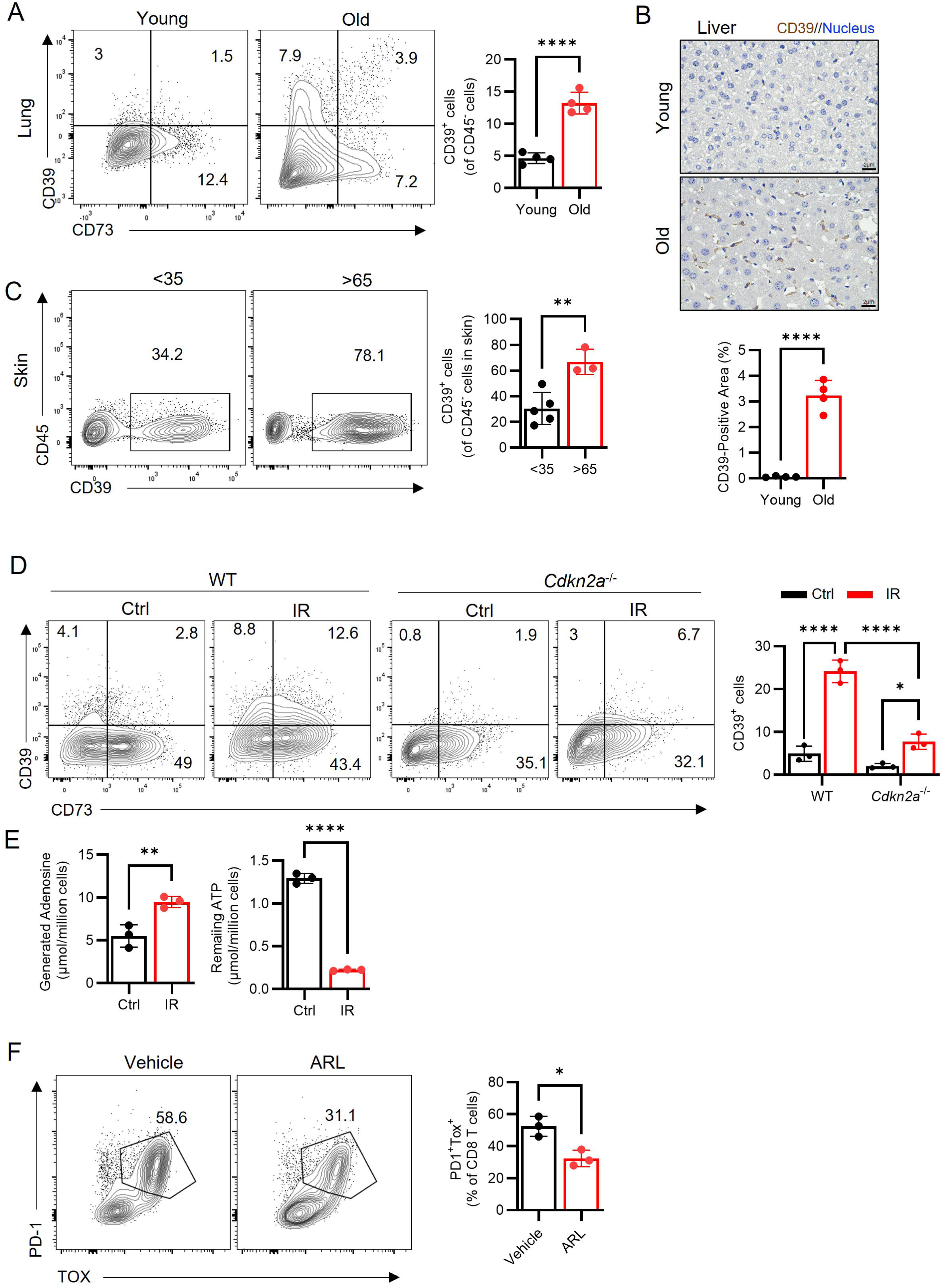
p16-dependent CD39 induction in senescent cells drives adenosine accumulation. (A) Expression of CD39 and CD73 on CD45^-^ cells in lung from young or old mice were analyzed by flow cytometry. (B) Immunohistology staining CD39 in liver from young and old mice. (C) Expression of CD39 on CD45^-^ cells in skins from young or old individuals were analyzed by flow cytometry. (D) MPFs isolated from WT or *Cdkn2a-*KO mice were subjected to irradiation (IR), one day later, CD39 expression was analyzed by flow cytometry. (E) MPFs were irradiated for one day, the medium was changed and 20 μm ATP was added, after one hour, the level of ATP and adenosine were measured. (F) NASH mice were generated as in Figure 1I, and ARL was administrated at last one month. The PD1^+^ TOX^+^ CD8 T cells in liver were analyzed by flow cytometry.

To determine whether cellular senescence can induce CD39 expression, we subjected primary mouse pulmonary fibroblasts (MPFs) to irradiation to trigger senescence. Irradiation robustly upregulated CD39 expression within one day (Fig. 3D) and maintained high expression for at least six days (data not shown). Similarly, CD39 was upregulated as well in serial passaging-induced senescent mouse embryonic fibroblasts (MEFs) (Fig. S6B). A large proportion of cells already expressed CD73 before irradiation or passaging, and its levels remained stable thereafter (Fig. 3D and fig. S6B), suggesting that CD39 represents the rate-limiting step for adenosine production in aged tissues.

We and other group previously showed that aged T cells also upregulate CD39 expression(*44, 48–50*), aged T cells are not typical senescent cells, they retain proliferative potential although exhibits some senescent-like phenotypes(*13–15*), implying that the mechanism driving CD39 expression in senescent cells differs from those in aged T cells. Because P16 is essential for both the onset and the phenotypic manifestations of cellular senescence, we examined its role and found that *Cdkn2a*-deficient MPFs (from homozygous *p16^Ink4a^*-luciferase reporter mice, as shown in fig. S6C) failed to upregulate CD39 following irradiation (Figure 3D), indicating that p16 is required for CD39 induction in senescent cells.

To determine whether irradiation-induced CD39 could function as an ecto-NTPDase, we incubated control and irradiated MPFs with ATP. Two hours later, ATP and adenosine levels were quantified. Irradiated MPFs degraded more ATP and generated more adenosine, in conjunction with CD73 activity (Fig. 3E). Consistent with these findings, treatment of aged mice with the CD39 inhibitor ARL67156 significantly reduced adenosine accumulation in multiple organs (Fig. 2B). Together, these results suggest that p16-dependent CD39 induction in senescent cells contributes to adenosine accumulation in aged tissues.

Given that elevated CD39 expression in senescent cells drives adenosine accumulation, thereby promoting Gzmk⁺ CD8 T cell differentiation, CD39 inhibition should reduce the generation of these cells. To test this, we re-established the NASH mouse model and treated animals with either vehicle or ARL67156. Remarkably, ARL67156 treatment markedly decreased the proportion of Gzmk⁺ CD8 T cells in the liver (Fig. 3F). Collectively, these findings demonstrate that p16-mediated CD39 induction in senescent cells elevates adenosine levels in aged tissues, which in turn drives the differentiation of Gzmk⁺ CD8 T cells.

### Eomes regulates Gzmk in response to adenosine and PD-L1 stimulation

Having established that Eomes is required for the differentiation of Gzmk^+^ CD8+ T cells (Fig. 2H), we next sought to determine whether Eomes directly dictates the *Gzmk* transcriptional program. Retroviral overexpression of Eomes in young CD8^+^ T cells was sufficient to markedly induce Gzmk expression (Fig. 4A). Conversely, CRISPR/Cas9-mediated ablation of *Eomes* in aged CD8 T cells significantly downregulated Gzmk production while concomitantly de-repressing Gzmb (Fig. 4B; fig. S7A–B). These gain- and loss-of-function studies demonstrate that Eomes acts as a master regulator that may directly drive *Gzmk* transcription while repressing the cytotoxic program.

**Figure 4.**
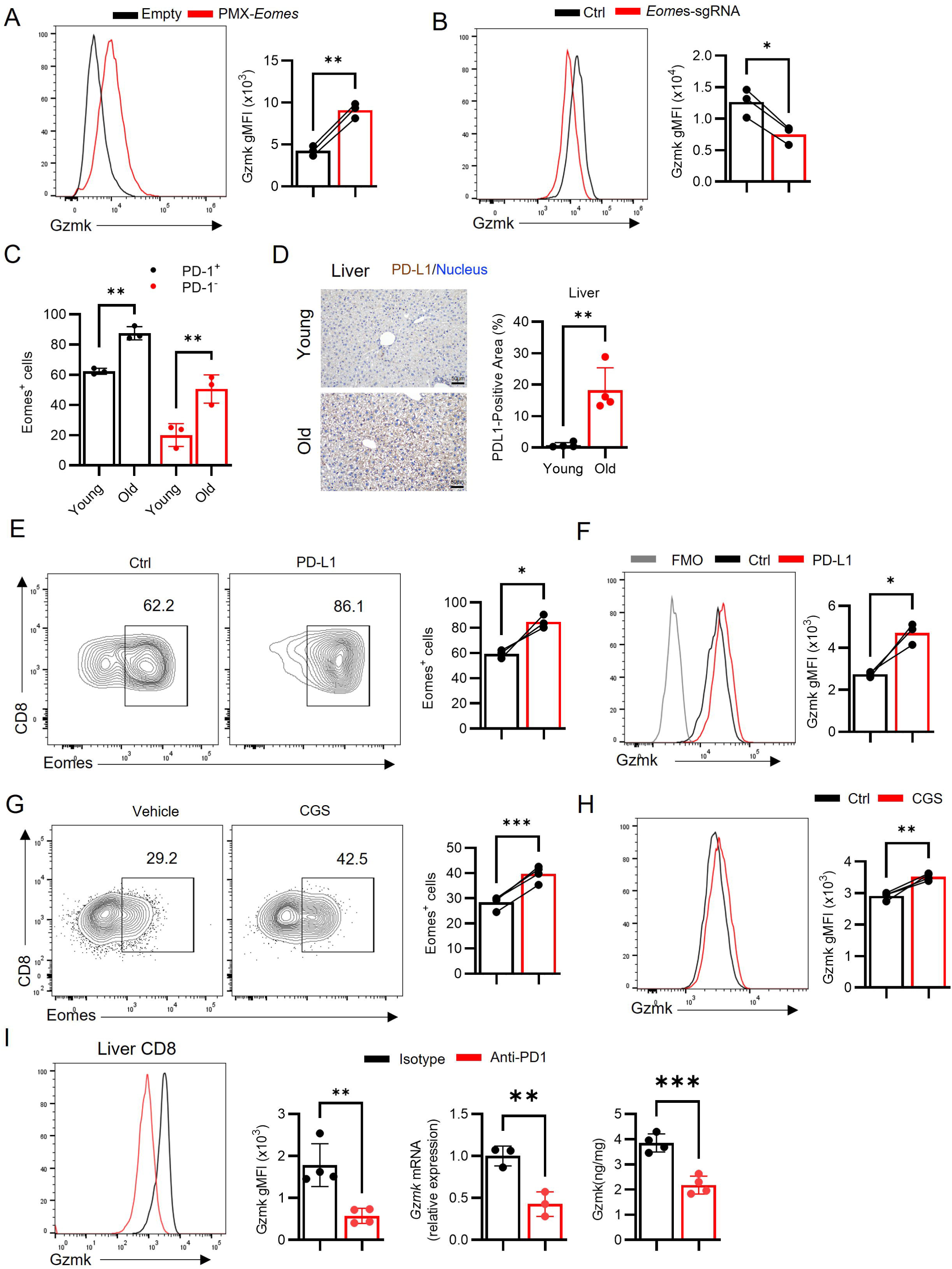
Eomes regulates Gzmk in response to adenosine and PD-L1 stimulation. (A) CD8 T cells isolated from spleen were transduced with empty or Eomes-retrovirus and cultured for three days, the expression of Gzmk were analyzed by flow cytometry. (B) Aged CD8 T cells purified from spleen were electroporated with control or *Eomes* sgRNA and Cas9 protein, then cultured for three days with anti-CD3/CD28 coated plate, the expression of Gzmk were analyzed by flow cytometry. (C) The expression of Eomes at PD1^-^ or PD-1^+^ CD8 T cells at livers from young or old mice were analyzed by flow cytometry. (D) Immunohistology staining PD-L1 in liver from young or old mice. (E-F) CD8 T cells isolated from old mice were cultured with anti-CD3/CD28 coated plate with or without PD-L1 (coating plate) for two days, Eomes expression (E), Gzmk expression (F) was determined by flow cytometry. (G-H) Young CD8 T cells isolated from spleen were cultured with anti-CD3/CD28 coated plate with or without CGS, Eomes expression (G), Gzmk expression (H) was determined by flow cytometry. (I) NASH mice were established as in Figure 1I, anti-PD1 antibody was administrated at last one month, Gzmk expression in CD8 T cells in liver was determined by flow cytometry, qPCR and ELISA.

We next investigated how the aging microenvironment sustains Eomes expression. Consistent with the accumulation of Gzmk^+^ cells, Eomes levels were significantly elevated in CD8 T cells resident in the aged spleen and liver (fig. S7C–D). Notably, this upregulation was not solely due to the expansion of the PD-1^+^ memory pool; PD-1^+^ CD8^+^ T cells from aged livers expressed significantly higher levels of Eomes than their phenotypic counterparts in young livers (Fig. 4C; fig S7E). This suggests that extrinsic cues within the aged niche actively maintain high Eomes expression in established Taa cells.

We hypothesized that two senescence-associated factors—PD-L1 and adenosine—converge to drive this Eomes-Gzmk axis. First, given the established link between senescence and PD-L1 upregulation(*22, 51*), we confirmed robust PD-L1 expression in aged liver (Fig. 4D). Functionally, PD-L1 ligation on aged CD8 T cells significantly potentiated both Eomes and Gzmk expression *in vitro* (Fig. 4E and F). Second, building on our finding that A2aR signaling is essential for Eomes induction *in vivo*, we cultured young CD8 T cells at present of CGS and activation of A2aR was able to upregulate Eomes and induce Gzmk (Fig. 4G and H).

Finally, we validated the functional requirement of the PD-1 axis *in vivo* using a NASH mouse model, which mimics the inflammatory stress of aging. Therapeutic blockade of PD-1 for two weeks markedly attenuated hepatic Gzmk expression (Fig. 4I). Collectively, these data define a feed-forward mechanism wherein senescence-associated adenosine and PD-L1 signaling converge on Eomes to lock CD8 T cells in a pathogenic Gzmk^+^ state.

### Genetic ablation of Gzmk extends lifespan and ameliorates age-related functional decline

To determine whether Gzmk^+^ CD8 T cells are causally involved in organismal aging, we carried out lifespan studies in *Gzmk*^-/-^ (pooled from male and female) and wild-type (WT) mice, which we observed until they were found dead or euthanized moribund. Strikingly, *Gzmk*^−/−^ mice exhibited a profound extension in longevity compared to WT controls, with the median lifespan increasing from 110 weeks in WT mice to 128 weeks in *Gzmk*^−/−^ mice (Fig. 5A).

**Figure 5.**
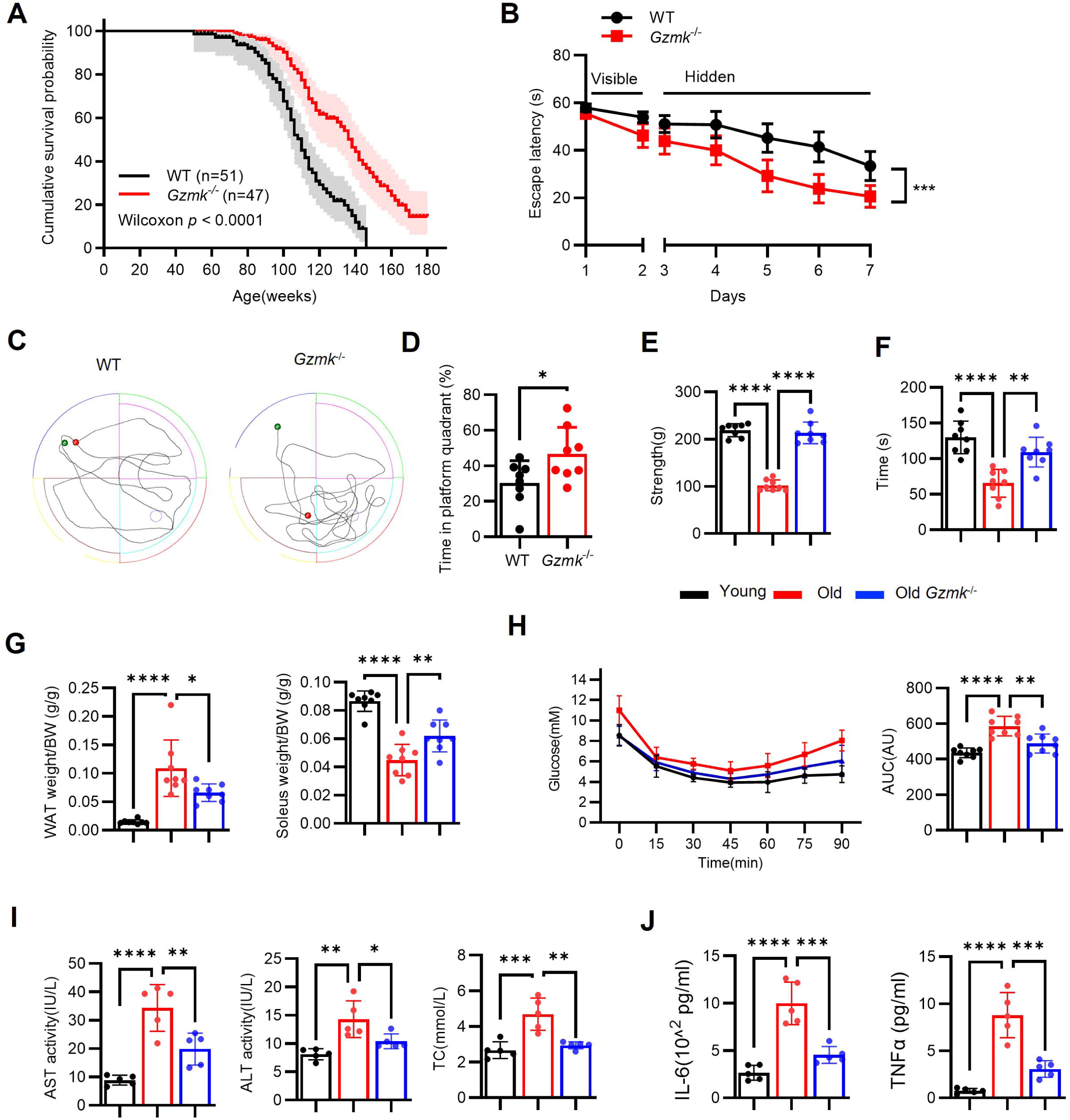
Genetic ablation of Gzmk extends lifespan and ameliorates age-related functional decline. (A) Analysis of survival probability of WT or *Gzmk*^-/-^ mice (from both female and male). (B-J) The analysis of male 19 months WT or *Gzmk*^-/-^ mice (male). (B-D) performance in Morris water maze. Grip strength (E), performance in rotarod (F). (G) The ration of WAT to BW and soleus to BW. (H) ITT were performed on indicated mice. The level of AST, ALT and TC (I), IL-6 and TNFα (J) in plasma were measured by ELISA.

Beyond lifespan extension, we investigated whether *Gzmk* deficiency improves healthspan. In behavioral assessments, old *Gzmk*^−/−^ mice displayed superior spatial learning and memory in the Morris water maze compared to WT controls (Fig. 5B-C), suggesting preserved cognitive function. Physical performance was similarly preserved: *Gzmk*^−/−^ mice exhibited enhanced muscle strength and physical endurance, as measured by grip strength and rotarod performance, respectively (Fig. 5E-F). Consistent with protection against age-associated physical functional decline, *Gzmk*^−/−^ mice maintained a higher soleus muscle-to-body weight ratio and a reduced white adipose tissue (WAT) burden (Fig. 5G). Metabolically, *Gzmk* deficiency attenuated age-related insulin resistance, as revealed by insulin tolerance tests (ITT). Moreover, Gzmk deletion significantly dampened the levels of aspartate aminotransferase (AST), alanine aminotransferase (ALT), and total cholesterol (TC), indicative of hepatic function, and pro-inflammatory cytokines in the serum (Fig. 5H and 5I). These findings demonstrate that *Gzmk* ablation broadly improves physiological function and delays the onset of frailty.

### CD8 T cell-derived Gzmk triggers systemic cellular senescent across organs

The progressive decline in organ function during aging is fundamentally driven by the systemic accumulation of senescent cells. To determine whether Gzmk directly contributes to this senescent burden, we evaluated senescence-associated β-galactosidase (SA-β-gal) activity across multiple tissues. Strikingly, SA-β-gal⁺ senescent cells were markedly diminished in the liver, brain, and adipose tissue of aged *Gzmk*⁻/⁻ mice compared to age-matched wild-type controls (Fig. 6A, B). This reduction in cellular senescence was accompanied by a profound protection against age-related structural deterioration. For instance, while natural aging typically induces pronounced senescence-associated hepatic steatosis, *Gzmk* deletion significantly attenuated lipid droplet accumulation (Fig. 6C). Furthermore, other classical histopathological hallmarks of aging, including alveolar enlargement in the lung(*52*), renal tubule dilation(*53*) and adipocyte hypertrophy(*54*) were significantly mitigated in *Gzmk*^−/−^ mice (Fig. 6C and fig. S8A). In parallel, the expression of key molecular senescence markers was substantially downregulated in the liver and brain of *Gzmk*-deficient animals (Fig. 6D and fig. S8B). To quantitatively and systemically monitor this senescent burden *in vivo*. we crossed *Gzmk*^−/−^ mice with *p16^Ink4a^*-luciferase reporter mice (fig. S6C). Bioluminescence imaging revealed that *Gzmk* deficiency markedly curtailed systemic luciferase activity in 15-month-old mice compared to age-matched controls, indicative of a significantly reduced burden of senescent cells (Fig. 6E). Collectively, these data establish that genetic ablation of *Gzmk* effectively restrains the systemic accumulation of senescent cells and preserves multi-organ structural integrity during aging.

**Figure 6.**
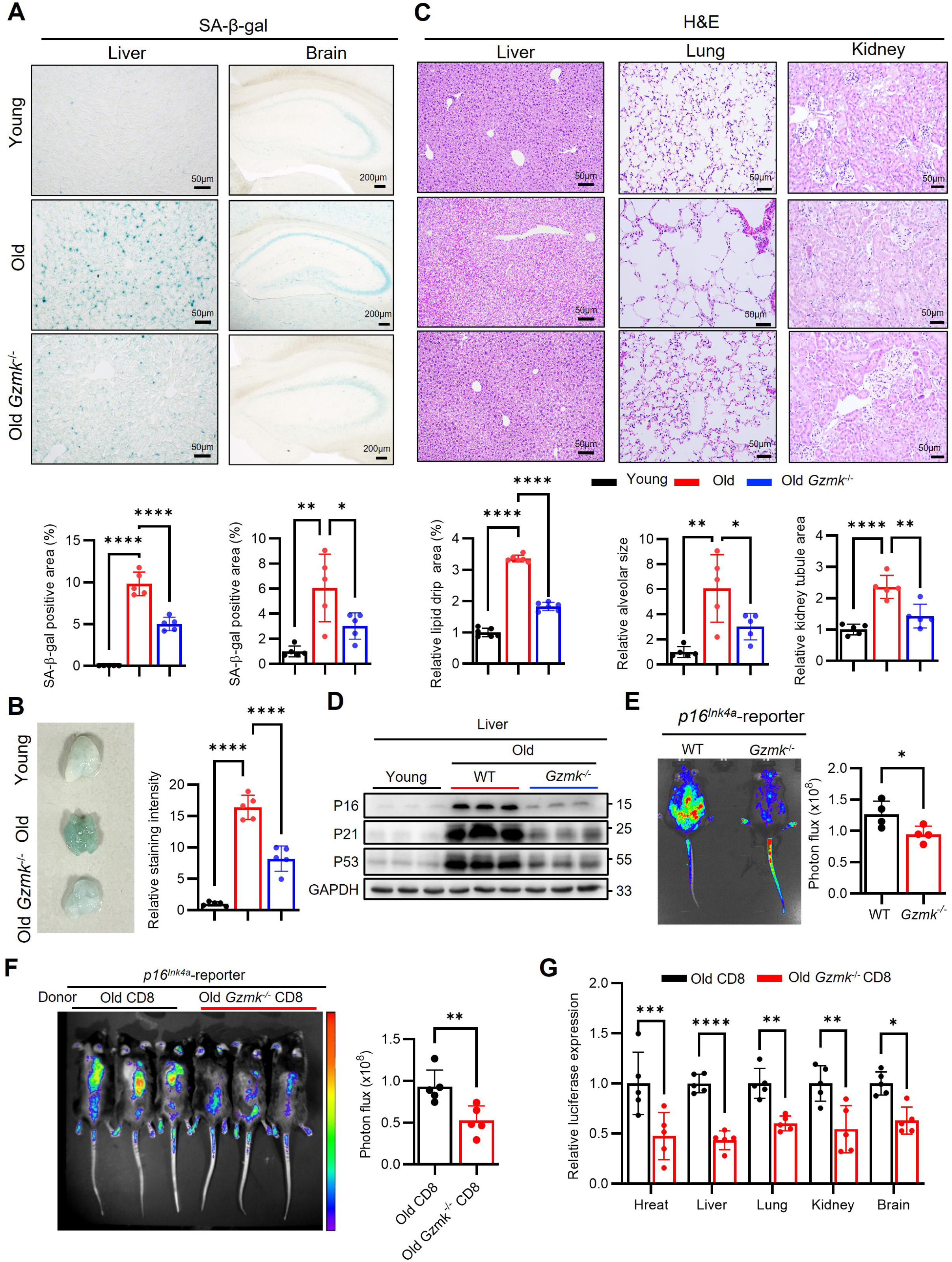
CD8 T cell-derived Gzmk triggers systemic cellular senescent across organs. (A-D) The analysis of male 19 months WT or *Gzmk*^-/-^ mice (male). (A) Staining of SA-β-gal in liver and hippocampus. (B) H&E staining in liver, lung and kidney. (C) SA-β-gal staining for WAT from indicated mice. (D) Western blot analysis of the expression of p16, p21 and p53 in liver. (E) The bioluminescence imaging of indicated mice (15 months, female) by injecting luciferase substrates. (F-G) CD8 T cells (4 x10^6^) from old or old *Gzmk^-/-^*mice were adoptively transferred into young *p16^Ink4a^*-luciferase reporter mice, one month later, the luciferase activity was measured (F), and the expression of the *p16*-driven luciferase reporter in indicated tissues were measured by RT-qPCR.

The diminished senescent burden observed in aged *Gzmk*⁻/⁻ mice could theoretically result from either impaired induction of senescence or enhanced immune-mediated clearance. Furthermore, because innate lymphocytes such as NK and NKT cells also express Gzmk, the specific contribution of CD8⁺ T cell-derived Gzmk remained to be definitively established. To address this, we isolated aged CD8⁺ T cells from wild-type or *Gzmk*⁻/⁻ donors and adoptively transferred them into young *p16^Ink4a^*-luciferase reporter mice One month post-transfer, young recipients of aged wild-type CD8⁺ T cells exhibited a strong induction of systemic bioluminescence, demonstrating that aged CD8⁺ T cells are intrinsically sufficient to drive premature *in vivo* (Fig. 6F). Strikingly, the adoptive transfer of aged *Gzmk*⁻/⁻ CD8⁺ T cells failed to elicit this robust systemic luciferase activity (Fig. 6F), This was corroborated by quantitative analysis, which showed markedly reduced induction of *Cdkn2a*, *Cdkn1a*, and luciferase transcripts in the tissues of recipients engrafted with *Gzmk*-deficient T cells (Fig. 6G and fig. S8C). Collectively, these results unequivocally establish CD8⁺ T cell-derived Gzmk not merely as a biomarker of aging, but as a potent and requisite paracrine driver of multi-organ cellular senescence.

### CD8 T cell-restricted *Gzmk* overexpression is sufficient to drive systemic inflammaging and premature tissue decline

To determine whether elevated CD8 T cell-derived Gzmk is sufficient to drive the systemic aging process, we engineered a transgenic mouse expressing murine *Gzmk* under control of *Cd8a* promoter and enhancer (fig. S9A). Among five independent founder lines screened, Line 1 (hereinafter referred to as *Gzmk*^TG^) was selected for subsequent studies due to its robust and specific expression profile (fig. S9B). We confirmed that *Gzmk*^TG^ mice exhibited constitutive, high-level Gzmk expression within CD8 T cells (fig. S9C), which effectively translated to elevated Gzmk protein levels in both circulating plasma and peripheral tissues (fig. S9D, E). Notably, *Gzmk^TG^* breeding pairs-irrespective of parental sex-yielded transgenic offspring at sub-Mendelian rations, hinting at early developmental or viability constraints imposed by systemic Gzmk overproduction (data not shown).

Strikingly, the targeted overexpression of Gzmk was sufficient to drive organ premature and a state of systemic inflammaging, evidenced by significantly elevated circulating the pro-inflammatory cytokines in young adult mice (Fig. 7A). Even on a normal diet, 8-month-old *Gzmk*^TG^ mice spontaneously developed severe hepatic steatosis, characterized by extensive parenchymal lipid droplet accumulation (Fig. 7C). This overt pathology was accompanied by a pronounced accumulation of senescent cells and impaired liver function (Fig. 7B, E). Beyond the liver, constitutive Gzmk expression induced age-related alveolar size expansion in lung (Fig. 7D). Lastly, we observed similar accelerated aging phenotypes in the brain of *Gzmk*^TG^ mice (Fig. 7E). Collectively, these *in vivo* gain-of-function studies demonstrate that restricted overexpression of Gzmk in CD8⁺ T cells is entirely sufficient to trigger widespread cellular senescence, inflammaging, and multi-organ structural decline, faithfully phenocopying the cardinal features of natural systemic aging.

**Figure 7.**
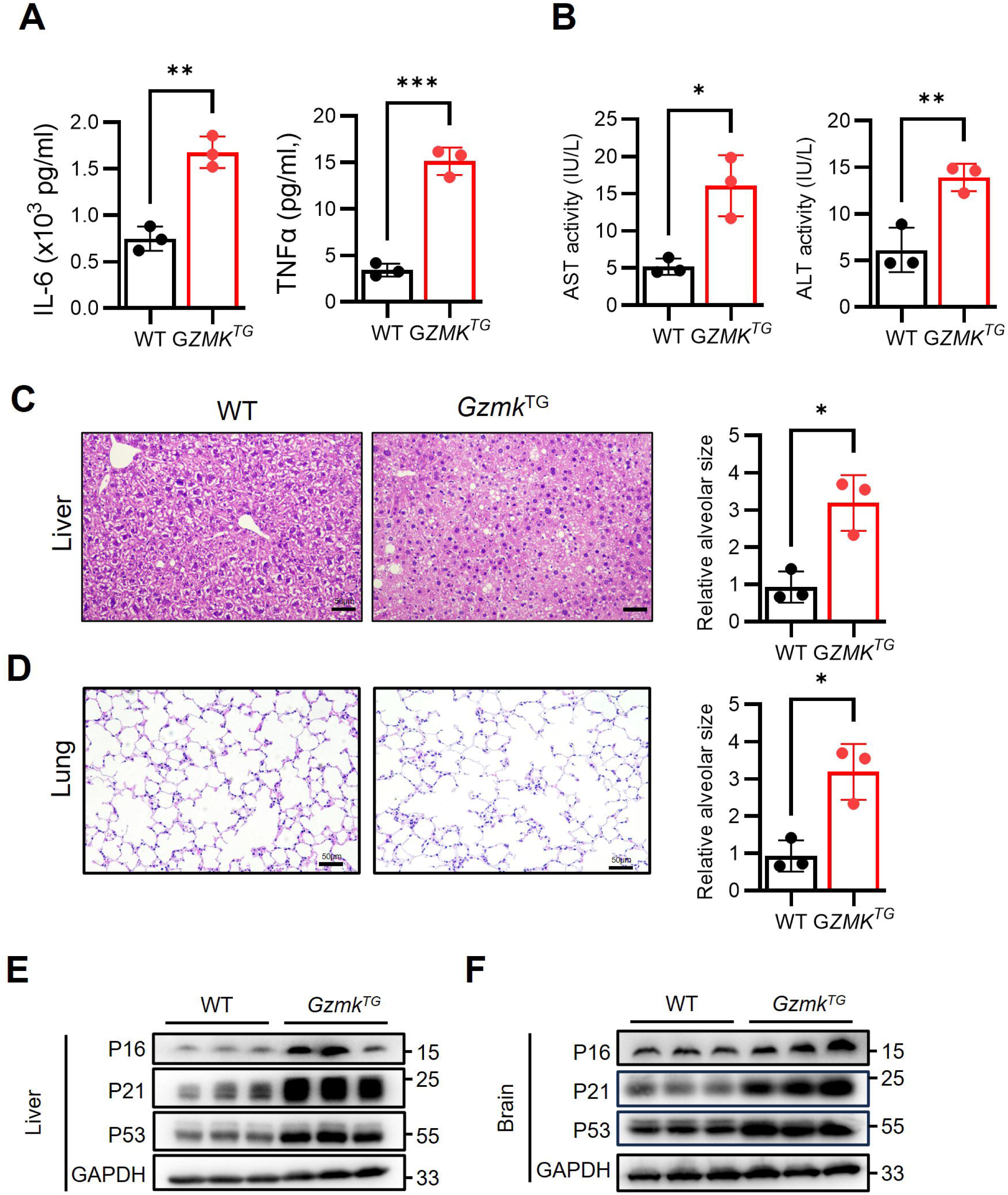
CD8 T cell-restricted *Gzmk* overexpression is sufficient to drive systemic inflammaging and premature tissue decline. 8 months WT or *Gzmk*^TG^ mice were analyzed. Level of IL6 and TNFα (A), AST and ALT (B) in plasma from WT or *Gzmk*^TG^ mice was measured by ELISA. H&E staining of liver (C), lung (D) from indicated mice. Western blot analysis of liver (E), brain (F) from 8 months WT or *Gzmk*^TG^ mice.

### Genetic ablation of *Gzmk* confers protection against senescence-associated NASH pathology

The pathogenic accumulation of senescent cells is a recognized driver of diverse age-related comorbidities, spanning from neurodegeneration to severe metabolic dysfunction. Having established that CD8⁺ T cell-derived Gzmk acts as a potent systemic inducer of cellular senescence, we next sought to determine whether *Gzmk* deletion could ameliorate tissue pathology in a defined disease context driven by localized senescence. We focused on nonalcoholic steatohepatitis (NASH), a progressive metabolic disorder intimately linked to hepatic senescence

Building on our earlier observation that pathogenic Gzmk⁺ CD8⁺ T cells specifically expand within the NASH-afflicted livers (Fig. 1G). We subjected WT and *Gzmk*^−/−^ mice to CDA-HFD to induce NASH. Comprehensive histological evaluation via Sirius Red and H&E staining revealed that *Gzmk* deficiency significantly attenuated hepatic fibrosis and lipid droplet accumulation (fig. 10A). This robust protection was accompanied by a downregulation of senescence-associated molecular markers (fig. 10B) and preserved liver function, as evidenced by restored plasma ALT and AST levels (fig. 10C).

Collectively, these findings underscore that targeted disruption of the Gzmk signaling axis not only mitigates the systemic physiological decline associated with natural aging but also confers potent therapeutic protection against senescence-driven, age-related metabolic pathologies.

### Extracellular Gzmk Drives Cellular Senescence and Inflammaging via a PAR1-MAPK Axis and Complement Activation

Building on our *in vivo* findings, we investigated whether extracellularly secreted Gzmk acts as a direct molecular effector of cellular senescence. Because Gzmk functions as a serine protease that cleaves extracellular substrates to propagate inflammation(*25, 27, 29, 55–58*), we treated human primary bladder fibroblasts (HBFs) with recombinant GZMK. Exposure to GZMK robustly induced senescence phenotypes accompanied by elevated SASP secretion (Fig. 8A, B), This potent pro-senescent and pro-inflammatory effect was broadly conserved, as GZMK triggered inflammatory cytokine production across multiple human cell lines (fig. S11A, B), and recombinant murine Gzmk elicited identical senescence programs in mouse embryonic fibroblasts (MEFs) (Fig. 8C, D).

**Figure 8.**
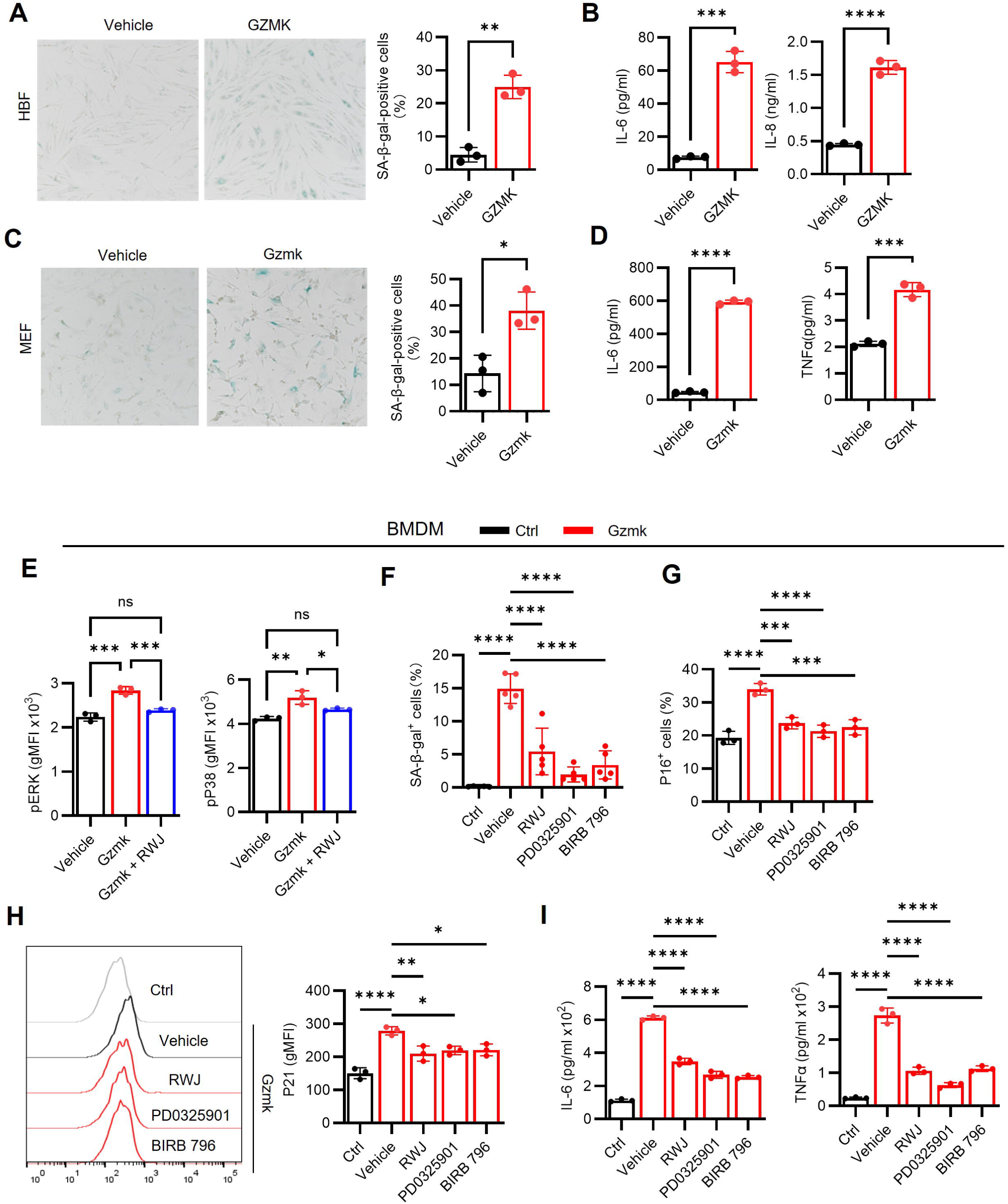
Extracellular Gzmk Drives Cellular Senescence and Inflammaging via a PAR1-MAPK Axis. (A, B) Primary human bladder fibroblasts were isolated from surgical excision from bladder cancer patients and subjected to GZMK treatment. (A) HBFs were treated with 100nM GZMK for 5 days, and SA-β-gal staining were performed. (B) HBFs were treated with 100nM GZMK in medium without FBS for one day, IL-6 and TNFα in supernatant were measured by ELISA. (C, D) MEFs were isolated and subjected to GZMK treatment as same with Figure 8A. (E-H) BMDMs were subjected to Gzmk or indicated inhibitors treatment. (E) BMDMs were stimulated with Gzmk for 30 minutes and phosphorylated ERK and p38 were determined by flow cytometry. (F) BMDMs were cultured with Gzmk or combined with indicated inhibitors for three days, SA-β-gal staining were performed. BMDMs were cultured with Gzmk or combined with indicated inhibitors for one day, expression of P16 (G) and P21 (H) were analyzed by flow cytometry, IL-6 and TNFα (I) in supernatant were measured by ELISA.

Since aged macrophages are potent drivers of systemic aging(*12*), and responsive to Gzmk-mediated activation(*26*). We next examined whether Gzmk triggers senescence in macrophage. To identify the receptor mediating these effects, we focused on protease-activated receptors (PARs), specifically PAR1, which was selectively and highly expressed on bone marrow-derived macrophages (BMDM) (Fig. S12A). Although PAR1 cleavage can trigger divergent downstream cascades depending on the biased proteases(*59, 60*), we discovered that Gzmk-mediated PAR1 activation specifically the phosphorylated ERK1/2 and p38 MAPK, but not NF-κB (Fig. 8E and fig. S12B). Given that sustained p38 and ERK1/2 signaling are established inducers of cellular senescence(*61–63*), we hypothesized that this axis drives the senescent phenotype mediated by Gzmk. Indeed, Gzmk treatment markedly induced senescence in BMDMs—evidenced by increased SA-β-gal^+^ cells, p16 expression, p21 upregulation, and production of IL-6 and TNFα. Importantly, pharmacological inhibition of PAR1, MEK, or p38 completely abrogated these Gzmk-induced senescence phenotypes (Fig. 8E-H and fig. S12C-F), establishing the PAR1-MAPK axis as the primary transducer of Gzmk signaling.

While this PAR1–MAPK axis dominates in isolated *in vitro* systems, the *in vivo* aged tissue microenvironment is characterized by markedly elevated levels of complement component 3 (fig. S13A, B). Because C3 is a known proteolytic target of Gzmk, we tested whether Gzmk-mediated complement activation serves as a parallel amplification mechanism. To test this, treatment of BMDMs with the cleavage product C3a elicited a mild increase in senescence markers (SA-β-gal, p16, p21) (fig. S13C-E), but significantly exacerbated SASP production (fig. S13F). These data suggest a dual mechanism wherein Gzmk primarily drives senescence induction via PAR1 while concurrently amplifying inflammation via complement activation.

### Pharmacological blockade of the A2aR–Gzmk axis reverses systemic inflammaging and multi-organ decline

To evaluate the translational and therapeutic potential of our findings, we investigated whether pharmacological targeting of the A2aR-Gzmk axis could recapitulate the anti-aging benefits of genetic ablation. We treated aged mice for one month with either the A2aR antagonist SCH 442416--to halt the induction of Gzmk^+^ T cell--or the serine protease inhibitor PPACK, to inhibit Gzmk enzymatic activity. Pharmacological blockade at either node of this axis significantly ameliorated age-associated systemic inflammation and hepatocellular injury, as evidenced by reduced plasma levels of TNFα, IL-6 AST and ALT (Fig. 9A, B). Consistently, treatment with SCH or PPACK markedly alleviated the burden of senescent cells and hepatic steatosis in the livers of aged mice (Fig. 9C, F). Similarly, both inhibitors ameliorated age-related alveolar changes in lung (Fig. 9D). Extending these benefits to the central nervous system, we found that inhibition of the A2aR-Gzmk axis attenuated hippocampal senescence (Fig. 9E, F).

**Figure 9.**
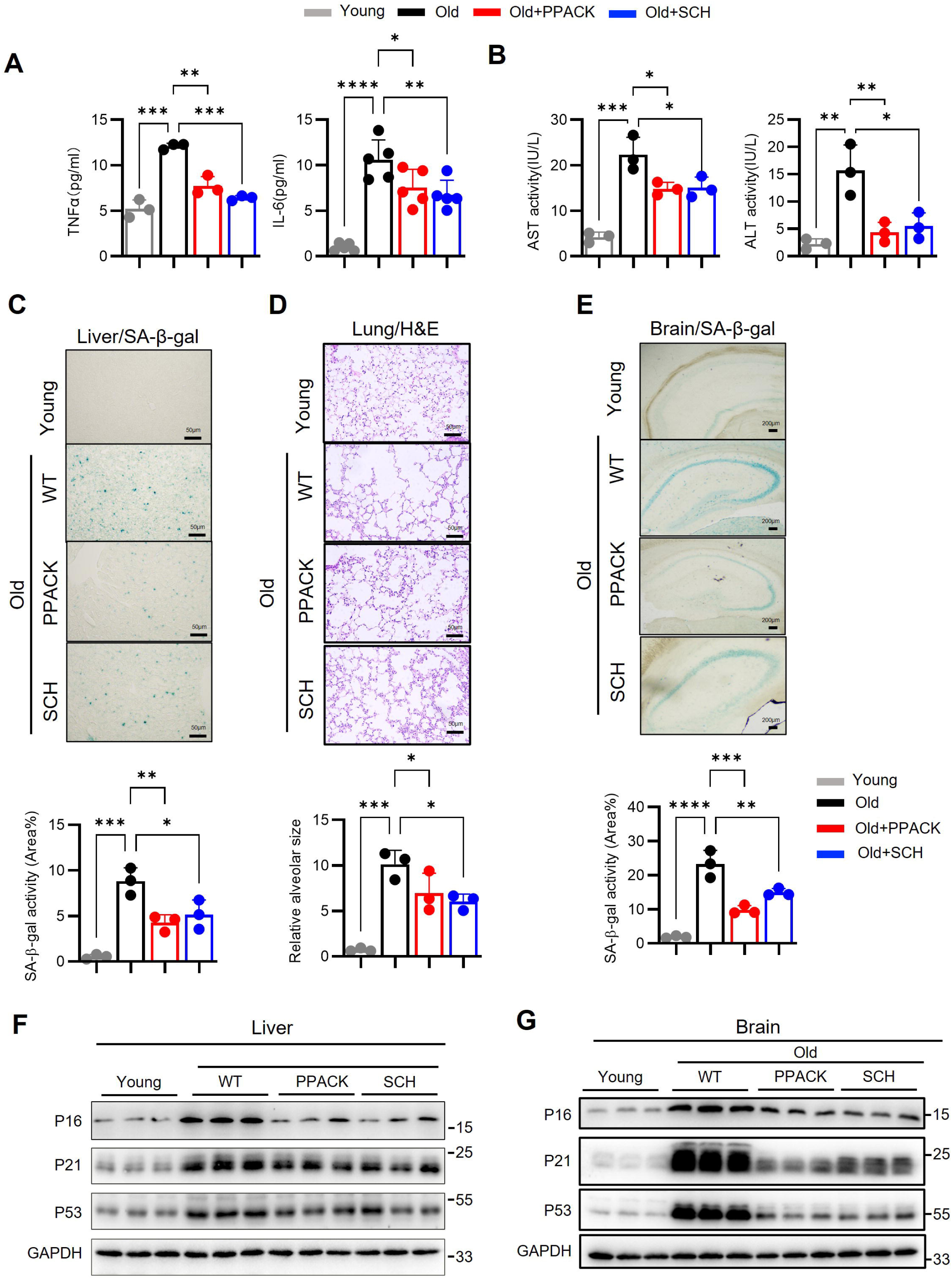
Inhibition of A2aR/Gzmk axis attenuates age-related phenotypes in aged mice. Old mice (19 months) were administrated with SCH or PPACK for one-month. The level of IL-6 and TNFα (A), ALT and AST (B) in plasma from indicated mice were determined by ELSIA. (C) Staining of SA-β-gal in liver. (D) H&E staining of lungs from indicated mice. (E) Staining of SA-β-gal in hippocampus from indicated mice. (F-G) Western blot analysis of P16, P21 and P53 at liver and brain from indicated mice.

Collectively, these preclinical data establish CD8⁺ T cell-derived Gzmk not merely as a consequence of aging, but as a systemic driver of organismal aging. These findings highlight the immense translational potential of therapeutically targeting the adenosine–Gzmk metabolic-immune axis—either by intercepting the upstream differentiation of pathogenic Gzmk⁺ T cells or by inhibiting their downstream proteolytic effectors—as a highly viable strategy to combat inflammaging and age-related multimorbidity.

## Discussion

T cell aging is canonically attributed to thymic involution or cumulative antigen/cytokine stimulation(*13–15, 64*), processes that deplete naïve CD8 T cells and skew the trajectory of memory populations(*65–67*), ultimately rendering older individuals susceptible to tumors and infections. Here, our study delineates a distinct mechanism: the aging tissue microenvironment actively “educates” T cells to become drivers of systemic aging. We identify a pathogenic axis wherein senescence-associated adenosine signaling reprograms CD8^+^ T cells into a stable Gzmk^+^ population via Eomes. These cells do not merely correlate with aging but actively propagate it through a Gzmk-PAR1-Complement feed-forward loop. This discovery unifies metabolic remodeling, immune dysfunction, and systemic inflammaging into a single causal framework, offering actionable targets—including the A2aR-Gzmk axis—to extend healthy lifespan.

The regulation of Gzmk presents a paradox: while initial TCR engagement is required for its differentiation(*32*), acute TCR stimulation actively represses Gzmk expression in human CD8 T cells(*27, 68*). This implies that once the Gzmk+ effector program is established, its maintenance and release become uncoupled from classical cognate antigen activation. This uncoupling is strongly supported by observations in chronic inflammatory conditions, wherein clonally expanded GZMK^+^ CD8 T cells, frequently recognize bystander antigens—such as those from Epstein-Barr virus (EBV)—rather than disease-driving autoantigens(*29*).

Mechanistically, we demonstrate that PD-1/PD-L1 and A2aR signaling converge to upregulate the transcription factor Eomes, thereby sustaining continuous Gzmk expression. It is well established that low-affinity TCR stimulation preferentially induces Eomes (*69, 70*). Because both PD-1 and A2aR function as canonical co-inhibitory receptors that dampen downstream TCR signaling cascades, we propose that the aged, adenosine- and PD-L1-rich microenvironment artificially mimics a “low-affinity” signaling state by restraining overall TCR signaling strength. This attenuated activation threshold perfectly optimizes and locks CD8 T cells into the Eomes–Gzmk pathogenic axis, thereby perpetuating inflammaging independent of acute antigen stimulation.

We identify Eomes as the master transcriptional regulator governing this pathogenic state. While Eomes is known to regulate virtual memory and exhausted CD8 T cell(*71–74*)—subsets that accumulate with age as well(*75, 76*)—our findings reveal a novel function: Eomes dictates the balance between cytotoxicity and inflammaging. Specifically, Eomes is required not only for Gzmk^+^ CD8 T cells differentiation but also for suppressing Gzmb in aged CD8 T cells. Indeed, Eomes ablation in aged CD8 T cells resulted in reciprocal upregulation of Gzmb (fig. 7B). This contrasts with its role in aged CD4 T cells, where Eomes drives a cytotoxic phenotype, which provides a protective role at aging(*21*). Whether adenosine similarly dictates Eomes induction in aged CD4⁺ T cells warrants further investigation. Our data suggest that in CD8 T cells, the aged microenvironment co-opts Eomes via A2aR signaling to repress “killing” machinery (Gzmb) in favor of “aging” machinery (Gzmk), thereby exacerbating systemic inflammation rather than clearing senescent cells.

This A2aR-mediated bias aligns with the classical immunosuppressive role of adenosine on T cells(*47, 77*), which likely impairs the immune clearance of senescent cells, further exacerbating the senescent burden. We demonstrate that the upregulation of CD39—the rate-limiting enzyme for adenosine generation—on senescent cells creates an adenosine-rich niche. This niche effectively switches the immune system from a surveillance state to inflammaging state by amplifying senescent cells accumulation. Consequently, although seemingly counterintuitive for an immunosuppressive pathway, A2aR blockade does not enhance inflammaging; rather, it profoundly attenuates it.This is line with PD-1 blockade, which ameliorates inflammaging by restoring the clearance of senescent cells(*22*). Our data show that PD-1 blockade also decreased the pro-aging Gzmk production by CD8 T cells. Therefore, similar to A2aR antagonism, PD-1 blockade favors the recalibration of CD8 T cells from an “aging” back to “killing” state. These findings prompt us to critically revisit the simplistic paradigm that combating aging relies solely on broadly suppressing inflammation.

While the ability of the aged microenvironment to imprint T cell dysfunction is recognized(*31, 32, 78*), the precise molecular driver has remained elusive. We identify extracellular adenosine as the critical metabolic cue instructing local Gzmk^+^ CD8 T cells differentiation in peripheral organs. Previous studies have demonstrated that TNFα accumulation and constitutive NF-κb activation in fibroblast establish an “inflammaging niche”, capable of driving the differentiation of this T cell subset(*31, 32*). Our findings integrate seamlessly with this paradigm, indicating that these inflammatory niches are fundamentally also adenosine-rich environments, driven by the senescence-associated upregulation of CD39.

However, our data and others suggest that metabolic instruction alone is insufficient for complete reprogramming. For instance, acute infection or immunization in aged, adenosine-rich hosts fails to induce Gzmk^+^ CD8 T cells(*32*). Similarly, our *in vitro* isolated A2aR stimulation (via CGS21680) modestly increased Gzmk expression but failed to generate the fully developed PD-1⁺ TOX⁺ phenotype (Fig. 4H and data not shown). Given that Gzmk^+^ CD8 T cells are generated locally within aged non-immune tissues, we propose a stringent “two-hit” requirement for their development during inflammaging: (1) atypical TCR priming orchestrated by senescent cells, acting synergistically with (2) sustained A2aR activation by adenosine. The precise molecular nature of this atypical priming—specifically how antigen presentation or co-stimulation by senescent cells diverges from that of their youthful counterparts—represents a critical frontier warranting further investigation.

In contrast to peripheral tissues, within the aged brain, adenosine levels are paradoxically diminished relative to young brains(*79*) (data not shown). Thus, the enrichment of Gzmk⁺ CD8 T cells in the aged brain is likely due to preferential migration rather than local differentiation. This migration model is supported by the fact high expression of the integrin CD49d on Gzmk⁺ CD8⁺ T cells (*24*) and the concurrent elevation of vascular cell adhesion molecule 1 (VCAM-1) on aged brain endothelial cells(*80*).

Downstream of this axis, we demonstrate that Gzmk drives systemic aging by acting on ubiquitously expressed substrates: PAR1 and complement components. By cleaving these targets, Gzmk initiates a proteolytic feed-forward loop that induces widespread cellular senescence. Importantly, the therapeutic implications of blocking this upstream metabolic driver are immediate. We show that pharmacological blockade of A2aR significantly reduces the Gzmk^+^ CD8 T cell burden and mitigates multi-organ aging phenotypes (Fig. 9). Furthermore, epidemiological evidence links habitual caffeine intake—a natural A2aR antagonist—to lower plasma Gzmk levels (Fig. 2). These findings bridge the gap between molecular mechanism and lifestyle intervention, suggesting that targeting the CD39-A2aR-Gzmk axis offers a viable strategy to extend healthy lifespan.

## Material and Methods

### Human subject study population

#### Human cohort about coffee consumption

The study population about coffee intake is from a human cohort designed originally for children obesity in Liaoning province. We included the parents’ samples with clear coffee intake information and no inflammatory diseases (ages 20 to 63, BMI 17.8 to 29.8 kg/m^2^, male 174, female 344). The study was approved by the Institutional Review Board of China Medical University.

#### Human Tissue Collection

All human tissue collection and subsequent experimental procedures were conducted in strict accordance with the principles of the Declaration of Helsinki. The study protocols were reviewed and approved by the Institutional Review Boards (IRBs) of China Medical University (including Shengjing Hospital and The First Hospital) and Shanghai Sixth People’s Hospital. Written informed consent was obtained from all participating patients or their legal representatives prior to surgical intervention.

#### Skin Specimens

Normal human skin specimens were obtained from macroscopically healthy surgical margins during the excision of benign cutaneous lesions. Fresh skin tissues utilized for flow cytometric analysis were procured from patients at Shengjing Hospital of China Medical University. For histological and multicolored immunohistochemical analyses, paraffin-embedded skin tissues—specifically sampled from sun-protected areas to exclude photoaging effects—were sourced from The First Hospital of China Medical University.

#### Renal and Bladder Tissues

Normal human renal and bladder tissues were acquired from patients undergoing radical nephrectomy for renal cell carcinoma (n=6, January–December 2025) or radical cystectomy for bladder cancer (January 2020–December 2025), respectively, at The First Hospital of China Medical University. To ensure the acquisition of strictly healthy tissues, a rigorous spatial and histological clearance protocol was employed. Immediately following complete resection, specimens were rinsed with ice-cold phosphate-buffered saline (PBS). Normal-appearing renal parenchyma or bladder mucosa, located at least 3 cm from the macroscopic tumor margin, was carefully dissected by an experienced urologist and divided into two portions. The first portion was subjected to independent evaluation by a board-certified pathologist to confirm the absolute absence of residual malignant cells. Upon histological clearance, the remaining tumor-free renal tissue was immediately processed for flow cytometric sorting and mass spectrometry imaging (MALDI-MSI). Correspondingly, the cleared normal bladder mucosa was utilized for the isolation of human primary bladder fibroblasts (HBFs).

#### Tonsil Specimens

Whole tonsil specimens were procured from six individuals undergoing routine surgical intervention for obstructive sleep apnea at Shanghai Sixth People’s Hospital. These macroscopically normal immune tissues were immediately processed for downstream flow cytometric and histological evaluations.

### Mice

C57BL/6J mice were purchased from GemPharmatech, Cd45.1 mice (Cat. NO. NM-KI-210226), and Cdkn2a-Luc-2A-tdTomato-2A-CreERT2 mice (Cat. NO. NM-KI-18039) were purchased from Shanghai Model Organisms Center, Inc.. Briefly, a Luc-2A-tdTomato-2A-CreERT2-WPRE-pA expression cassette was inserted into the Cdkn2a locus via homology-directed repair (HDR). The knock-in was targeted to exon 2 of the Cdkn2a-201 transcript, immediately downstream of the codon encoding the 62nd amino acid. 5xFAD (034840-JAX) were originally from the Jackson Laboratory. Gzmk-KO mice (Strain S-KO-02387) were purchased from Cyagen. *Gzmk* transgenic mice were generated by inserting a customized genetic construct in which the Cd8a promoter drives the expression of a Kozak-Mouse Gzmk coding sequence fused to a P2A-EGFP cassette. The targeting vector consisted of the E8I enhancer, Cd8a promoter, Kozak motif, Gzmk CDS, the self-cleaving P2A peptide, an EGFP reporter, the WPRE regulatory element, and a BGH polyadenylation signal, together with a kanamycin resistance cassette used for selection during vector construction (generated by Cyagen).

Mice were housed 2–5 to a cage at an ambient temperature of 23–25 °C in a humidity-controlled room and were maintained on a 12-h light/dark cycle (08:00 to 20:00 light on) with standard food (CA-1, CLEA) and water provided ad libitum and with access to environmental enrichment. The experimental procedures were conducted according to the Chinese regulations involving animal protection and approved by the Animal Ethics Committee of the China Medical University.

Both male and female mice were used in all experiments, unless otherwise specified. Young mice are 3-4 months, and old mice are 18-24 months. For each experiment, at least three animals were included in each group, and data were pooled from 2–3 independent experiments. Sex- and age-matched animals of the indicated genotypes were randomly assigned to different groups in each experiment.

### Mouse model and treatment

For induction of the NASH model, 6-8 weeks mice were fed an L-amino-acid–defined high-fat diet containing 60Lkcal% fat with 0.1% methionine and no added choline (CDA-HFD; Huafukang, Beijing, China) for a total of 10 weeks.

For induction of the pulmonary aging model, a single dose of bleomycin (BLM; Cat#: HY-17565A; MCE, Shanghai, China) was administered intratracheally using a MicroSprayer aerosolizer (Model YAN 30012; Yuyan Instruments, Shanghai, China). Briefly, 50 µL of BLM solution at a concentration of 2.5 mg/mL was loaded into the MicroSprayer and delivered into the trachea of anesthetized mice.

For pharmacological intervention, ARL67156 (2 mg/kg; MCE), SCH (1 mg/kg; MCE), PPACK (62.5ug per mouse per time, HY-122542B, MCE), in a volume of 200 μL were administered every other day by intraperitoneally injection for indicated time. All mice were perfused and tissues were collected 24 hours after the final injection.

### Primary murine cell cultures

All data points referring to murine samples represent distinct mice. Total CD8 T cells were isolated from spleen using using EasySep kits (EasySep Mouse CD8+ T Cell Isolation Kit #19853). The purity of sorted cells was assessed by flow cytometry Only samples with purity >90% were used for subsequent experiments. For invitro culture experiments, cells were activated with plate-bound anti-CD3/anti-CD28 antibodies (both 5 μg/mL, Biolegend) for 2 or 3 days. When indicated, CGS21680 (5 nM, MCE) was added at the beginning of culture. For PD-L1 stimulation, plate-bound PD-L1 was used (chimeric mouse PD-L1/Fc, 3 μg ml−1; 1019-B7-100 R&D).

For overexpression, mouse Eomes CDS was inserted into PMX vector(*81*). Virion preparation was generated in the PLAT-E cell line, retrovirus-containing supernatant was collected after 72 hours. For transduction, CD8 T cells were purified young or old mice splenocytes by negative selection (STEMCELL Technologies) and stimulated in 48-well plates precoated with 5 μg/mL anti-CD3 and anti-CD28. Twenty-four hours after stimulation, cells were transduced with retrovirus-containing supernatants supplemented with 50 μM BME and 8 μg/mL polybrene (MilliporeSigma) by centrifugation for 90 minutes at 1500 *g* at 32°C.

For Cas9 mediated knock-out, sgRNAs were pre-assembled in vitro with recombinant Cas9 protein to form ribonucleoprotein (RNP). CD8 T cells from aged mice using negative selection, and RNP complex was electroporated into CD8 T cells using the Lonza 4D-Nucleofector™ System (Lonza) in combination with the P3 Primary Cell 4D-Nucleofector™ X Kit S (Lonza, Cat. No. V4XP-3032) according to the manufacturer’s instructions. The electroporated CD8 T cells were activated with plate-bound anti-CD3/CD28 antibodies for 2 days.

### Isolation and culture of primary bone marrow macrophages

The hind limbs were removed from mice; sterile PBS was repeatedly injected into the bone marrow cavity to flush out the bone marrow. After thoroughly flushing the bone marrow, the suspension was filtered and centrifuged at 1600 rpm for 5 minutes. The supernatant was discarded; the pellet was resuspended in 1 mL of RPMI 1640 medium supplemented with 15 ng/mL M-CSF and plated into culture wells. The medium was changed on day 3 and day 5. On day 6, cells were harvested and stained with CD11b and F4/80 for flow cytometry to assess purity. Afterward, cells were replated for subsequent drug treatment and stimulation. Recombinant Gzmk (100 nM, MBS2029386, MyBioSource), PAR inhibitor (10 μM, RWJ-56110, MCE), MEK inhibitor (10μM, PD0325901, MCE), or p38 inhibitor (10μM, BIRB 796, MCE) was used.

### Isolation and culture of primary microglia

Neonatal mice within 48 hours of birth were used. The cerebral cortex was dissected and placed in dissection solution (HBSS supplemented with 1% Penicillin-Streptomycin (P/S)). After removing the meninges under a stereomicroscope, the tissue was minced into small pieces in a centrifuge tube containing 1 mL of HBSS with 1% Fetal Bovine Serum (FBS) and 1% P/S. Subsequently, 1 mL of trypsin was added, and the tube was incubated for 15 minutes at 37°C in a CO₂ incubator, with gentle shaking every 5 minutes. Digestion was terminated by adding 2 mL of culture medium, followed by centrifugation. The supernatant was discarded, and the fluffy pellet was collected, resuspended in 1 mL of culture medium by gentle pipetting, and set aside. Dishes were pre-coated with 10 µg/mL Poly-D-lysine (PDL) and incubated at 37°C for at least 4 hours. After incubation, the dishes were washed three times with ddH₂O. The resuspended cell suspension was then plated into the coated dishes containing 3 mL of DMEM/F12 culture medium. The medium was completely replaced on the following day (day 2), followed by half-medium changes every 3-4 days thereafter, until the dishes were confluent with microglia. The supernatant containing loosely adherent microglia was gently collected and cultured. Following the addition of 1 mL of trypsin, the dish was gently tapped to detach the remaining microglia, which was monitored under a microscope. The digestion was stopped by adding 1 mL of culture medium. All supernatant was collected, centrifuged, and the cell pellet was assessed for purity. After confirmation, the cells were replated for subsequent drug treatment or stimulation experiments.

### Isolation and culture of Mouse Embryonic Fibroblasts (MEFs)

MEFs were isolated from E12.5–E13.5 mouse embryos. After removal of the head and visceral tissues, the remaining trunk tissue was minced and digested with trypsin at 37°C for no longer than 30 minutes until the tissue was fully dissociated without visible large fragments. Following termination of digestion, a single-cell suspension was prepared and cultured in DMEM supplemented with 10% fetal bovine serum (FBS) and 100 U/mL penicillin/streptomycin. The medium was replaced after 24 hours to remove non-adherent cells.

### Isolation and culture of Mouse Pulmonary Fibroblasts (MPFs)

MPFs were isolated from lung tissues of 6–8-week-old WT or *Cdkn2a*-KO mice. The lungs were minced into small fragments and digested with collagenase type II (2 mg/mL) at 37°C for 45 minutes to obtain a single-cell suspension. After digestion, the suspension was filtered through a cell strainer and centrifuged to collect cells. Cells were cultured in DMEM supplemented with 10% fetal bovine serum (FBS) and 100 U/mL penicillin/streptomycin. The medium was replaced 24 hours after seeding to remove non-adherent cells. Cells were subsequently passaged every 2–3 days.

### Isolation and culture of human bladder tumor–derived fibroblasts (HBFs)

Fresh bladder tumor tissues (0.2–1.0 g) were washed twice with phosphate-buffered saline (PBS) and minced into small fragments. Tissue dissociation was performed using a Human Tumor Tissue Gentle Dissociation Kit (Cat# DHTEH-2505, RWD) according to the manufacturer’s instructions, assisted by a single-cell suspension dissociator (DSC-410, RWD). After digestion, cell suspensions were filtered through a 70-μm cell strainer and centrifuged at 500 × g for 5 min to collect cell pellets. Pellets were resuspended in RPMI medium (Cat# 6125437, Gibco) supplemented with 10% fetal bovine serum (FBS) and 100 U/mL penicillin–streptomycin and plated in 10-cm culture dishes. Cells were maintained under standard culture conditions, and the medium was replaced every 3 days.

### Adoptive transfer

For A2aR or Eomes knock-out experiments, CD8 T cells isolated from CD45.1 Cas9-expressing mice were electroporated with indicated sgRNA (CUCGCUCUUCCAGUACCCGG for Eomes; GAAUUCCACUCCGGUGAGCC for A2aR), and were rested for three hours, then were transferred into old mice (one million per mouse) via tail vein injection. Splenocytes were analyzed one month later. For brain migration experiments, PD1^+^ CD44^+^ or PD1^-^ CD44^+^ CD8 T cells from old CD45.1 mice (lymph nodes + spleen pooled from 3 mice) were sorted were sorted on a BD FACSAria III, and were transferred to young or old receipts (one million per mouse), spleen and brain were analyzed after two weeks.

### Flow Cytometry

Cells were stained with Fixable Viability Dye eFluor™ 506 (Thermo Fisher, cat# 65-0866-18) and blocked with mouse or human TruStain FcX (BioLegend, cat# 101319 or 422302). Surface markers were stained by incubation with fluorochrome-conjugated antibodies for 30□min at 4°C in the dark.

For intracellular cytokine staining (Gzmk and Gzmb), cells were stimulated for 4□h at 37°C under 5% CO₂ in RPMI-1640 medium containing 10% FBS, 100□U/mL penicillin/streptomycin, 50□μM β-mercaptoethanol, 2.5□ng/mL PMA (Sigma, P8139), 1□μM ionomycin (Sigma, I3909), and GolgiStop (BD Biosciences). Cells were then fixed and permeabilized using the Fixation/Permeabilization Solution Kit (BD, cat# 554715) or the True-Nuclear Transcription Factor Buffer Set (BioLegend, cat# 424401) according to the manufacturer’s instructions, followed by intracellular antibody staining.

Samples were acquired on a FACSCelesta flow cytometer (BD Biosciences). Data were collected with FACSDiva software and analyzed using FlowJo v10.8.1.

The following antibodies were used:

CD45-PerCP-Cy5.5(clone 30-F11, BioLegend, 103131),CD8-APC-Cy7 (BioLegend, 100714, clone 53-6.7),CD4-AF700 (BioLegend, 100430, clone GK1.5),PD-1-PE (Invitrogen, 11-9969-42, clone MIH4),PD-1-APC (BioLegend, 135210, clone 29F.1A12),CD39-PE-Cy7 (BioLegend, 143806, clone Duha59),CD73-APC (BioLegend, 127209, clone TY/11.8),TCR β chain-Pacific Blue (BioLegend, 109226, clone H57-597),TOX/TOX2-AF488 (CST, 44682S),PD-L1-FITC (Elabscience, E-AB-F1132UC, clone 10F.9G2),CD44-PerCP-Cy5.5 (BioLegend, 103032, clone 1M7), CD45.1-AF700 (BioLegend, 110723, clone A20),Eomes (BD, 566749, clone X4-83).

### Isolation of single-cells from human tissues

For human skin, fresh full-thickness human skin was cut into ∼1-mm-thick tissue fragments and incubated with Dispase® II (0.2%; Sigma-Aldrich) at 37□°C for 2 h to separate the epidermis and dermis. The isolated epidermis was further digested with trypsin–EDTA at 37□°C for 30 min. Dermal single-cell suspensions were generated by incubating the dermis at 37□°C for 2 h under continuous agitation in digestion buffer containing collagenase type I (0.2% final; Invitrogen) and DNase I (30 IU/mL; Sigma).Following digestion, cell suspensions were collected, passed through a 70-μm cell strainer, and centrifuged at 500 × g for 5 min. Cell pellets were resuspended in RPMI supplemented with 10% fetal bovine serum (FBS), penicillin–streptomycin (100 U/mL), and L-glutamine, and cultured overnight under standard conditions. Cells were collected the next day for subsequent flow cytometry staining and analysis. For human kidney, fresh human kidney tissue was transported on ice in RPMI and processed as soon as possible. Visible fat, blood clots, and necrotic areas were removed, and the tissue was minced into ∼1–2 mm fragments using sterile blades. Tissue fragments were incubated in RPMI containing collagenase D (1 mg/mL final) and DNase I (20 μg/mL) at 37°C in a shaking incubator for 30 min. After digestion, the suspension was filtered through a 70-μm cell strainer and centrifuged at 500 × g for 5 min at 4°C to collect the cell pellet. Cells were resuspended in PBS and stained with the indicated antibodies prior to flow cytometric acquisition. For tonsil, tonsil tissues were minced into ∼5 mm³ fragments and dissociated into single-cell suspensions by mechanical disruption. The suspensions were then filtered through a 70-μm strainer.

### Isolation of mononuclear cells from mouse tissues

Mice were anesthetized with ether (Sigma) and perfused transcardially with PBS to remove circulating blood cells. Liver and brain tissues were finely minced with sterile scalpels and digested at 37°C for 30 min in RPMI medium (Cat#: 6125437, Gibco) containing 1 mg/mL Collagenase D (Cat#: 11088858001, Roche) and 50 μg/mL DNase I (Cat#: 10104159001, Sigma) using a single-cell suspension dissociator (DSC-410, RWD). Following digestion, all samples were passed through a 70-μm cell strainer and centrifuged at 500 g for 10 min to collect cell pellets. The supernatant was discarded, and pellets were resuspended using the High Efficiency Debris Removal Kit (Cat#: DHDR-5006, RWD). According to the manufacturer’s instructions, the appropriate volume of pre-chilled PBS was gently layered along the tube wall, followed by centrifugation at 2700 g for 20 min at 4°C with slow acceleration and deceleration. After centrifugation, the upper two layers were completely removed, and cells in the bottom layer were collected. The pellet was resuspended in ice-cold PBS and centrifuged again at 1000 g for 10 min at 4°C. The supernatant was removed, and the final cell pellet was resuspended in PBS to the desired volume for downstream applications.

Lung tissues were harvested, finely minced, and digested in HBSS supplemented with 1 mg/mL collagenase IV (Gibco, 17104019) and 20 μg/mL DNase I (Sigma, 10104159001) at 37°C for 2 hours with gentle shaking. The resulting cell suspension was passed through a 70-μm strainer and centrifuged at 2000 rpm for 5 minutes at 4°C. Red blood cells were removed using ACK lysis buffer, and viable cells were subsequently counted for downstream analyses.

### Quantification of extracellular ATP and adenosine

Adenosine levels in tissue lysates and culture supernatants were measured using an adenosine assay kit (MET-5090, Cell Biolabs). ATP concentrations were determined using a chemiluminescent ATP assay kit (E-BC-F300, Elabscience). All assays were performed according to the manufacturers’ instructions.

### Morris Water Maze Test

The Morris water maze (MWM) test was performed in a circular stainless-steel pool (120 cm in diameter, 50 cm in depth). The pool was filled with tap water, which was rendered opaque by the addition of milk to conceal the submerged platform. Water temperature was maintained at 23–25 °C throughout the experiment. Distinct geometric cues were placed on the walls surrounding the pool in each of the four quadrants as spatial references. All trials were recorded and analyzed using a video tracking system (Panlab, Smart 3.0).

A transparent circular escape platform (10 cm in diameter) was placed in the center of the target quadrant (southwest quadrant). Training lasted for 7 consecutive days and consisted of two phases: the visible platform phase (days 1–2) and the hidden platform phase (days 3–7). During the visible platform phase, the platform was positioned 1 cm above the water surface, whereas during the hidden platform phase, it was submerged 1 cm below the water surface. Each mouse underwent 3 trials per day, starting from three different quadrants in a pseudo-random order. In each trial, the mouse was allowed 60 s to locate the platform. If the mouse failed to find it within the allotted time, it was gently guided onto the platform and remained there for 30 s before being returned to its home cage.On day 8, a probe test was conducted to assess spatial memory retention. The platform was removed from the pool. Each mouse was released from the quadrant opposite the former platform location (northeast quadrant) and allowed to swim freely for 60 s. The number of crossings over the previous platform location and the time spent in the target quadrant were recorded and analyzed.

### Rotarod Test

#### 1. Training Phase

On the day before testing, mice were acclimated to the testing room for 30 minutes. For training, mice were first placed on a stationary rod for 5 min, then exposed to a constant low-speed rotation (3 rpm for 1 min). Subsequently, the speed was gradually increased up to 30 rpm over a 5–10 min training session in accelerating mode. Mice that fell were promptly returned to the rod.

#### 2. Test Procedure

On the test day, mice were again acclimated for 30 minutes before the trial. Each mouse was gently placed on the stationary rod facing away from the direction of rotation. The trial began with the rod rotating at 3 rpm, accelerating linearly to 30 rpm over 5 min (300 s). The latency to fall was recorded automatically. The test was terminated if the mouse fell, clung to the rod for more than 2 s, or completed the full 300s period. Each mouse performed three trials per day with a minimum inter-trial interval of 15 min. Testing was repeated for three consecutive days, and the average latency from the third day was used for analysis.

### Grip Strength Test

Grip strength was measured in mice using a digital grip strength meter. Mice were acclimated to the testing room for 30 min. For forelimb strength measurement, each mouse was gently held by the tail and allowed to grip a metal grid with its forepaws only. The mouse was then pulled steadily backward until grip release, and peak force was recorded. Each mouse underwent five consecutive trials with 30s intervals. The maximum value from the five trials was used for analysis. Data were normalized to body weight and analyzed using unpaired t-test.

### ITT test

For ITT, Mice were fasted for 6 h prior to baseline blood glucose measurement were injected intraperitoneally with recombinant human insulin(NovoMix 30 penfill,novo nordisk) at 0.75 mU per g body weight. Blood glucose concentrations were then measured at 15, 30, 60, 75 and 90 min after insulin administration. Both glucose and insulin were diluted in sterile PBS. respectively. Blood was collected via tail snip and yuwell blood glucometer was used for blood glucose measurements.

### Enzyme-linked immunosorbent assays (ELISA)

Plasma activities of alanine aminotransferase (ALT/GPT) and aspartate aminotransferase (AST/GOT) in mice were measured using ALT Activity Assay Kit (E-BC-K235-M, Elabscience) and AST Activity Assay Kit (E-BC-K236-M, Elabscience), respectively. Total cholesterol (TC) levels in mouse plasma were determined using a colorimetric assay kit (E-BC-K109-M, Elabscience). Plasma concentrations of IL-6 and TNF-α in mice were quantified using mouse micro-sample IL-6 ELISA kit (E-MSEL-M0001, Elabscience) and mouse micro-sample TNF-α ELISA kit (E-MSEL-M0002, Elabscience), respectively. Mouse granzyme K (GZMK) levels were measured using a mouse GZMK ELISA kit (JL31725, Jianglai Biotechnology). Human serum GZMK levels were determined using a human GZMK ELISA kit (JL31723, Jianglai Biotechnology).

For tissues, tissue samples were weighed and homogenized in ice-cold PBS at a weight-to-volume ratio of 1:9 (1 g tissue in 9 mL PBS). Homogenates were centrifuged at 12,000 ×g for 15 min at 4°C, and the supernatants were collected as tissue lysates for subsequent.

For soluble and insoluble Aβ, tissue samples were homogenized in TBS buffer at a ratio of 10 mg tissue per 100 μL buffer and centrifuged at 12,000 ×g for 1 h at 4°C. The supernatants were collected as the soluble Aβ fraction. The resulting pellets were resuspended in lysis buffer containing 1% Triton X-100, thoroughly pipetted, and subjected to ultrasonic disruption, followed by incubation at room temperature for 2 h. Samples were then centrifuged at 12,000 ×g for 40 min at 4°C, and the supernatants were collected as the insoluble Aβ fraction. Tissue Aβ42 levels were quantified using a Human Aβ1-42 (Amyloid Beta 1-42) ELISA Kit.

### SA-**β**-gal assay

SA-β-gal activity was assessed with a senescence β-galactosidase staining kit (9860S, CST), according to manufacturer’s protocol. For cells, indicated cells were washed three times with PBS and fixed with 10X Fixative solution at room temperature for 15 min. After washing with PBS, fibroblast cells were stained with 1 ml of freshly prepared β-gal staining solution at 37°C overnight, BMDM were stained with 1 ml of freshly prepared β-gal staining solution for 8 hours at 37°C. The cells were washed twice with PBS and overlaid with PBS. Images were captured under a microscope (Nikon). Blue-stained cells were indicated for cellular senescence. For tissues staining, frozen liver section and brain section were stained with prepared β-gal staining solution at 37°C overnight after washing with PBS. Images were captured under a microscope (Nikon). White adipose tissue fixed in 4% paraformaldehyde was rinsed and immersed in β-Gal staining solution at 37°C overnight. Staining intensity was assessed macroscopically.

### Imaging MS

The tissues were removed immediately frozen in liquid nitrogen and stored at −80°C until use. The tissues were sliced at a thickness of 10 μm with a cryostat (Leica CM1950, Nussloch, Germany) at −23°C (cryochamber temperature) and −16°C (specimen cooling temperature). The slices were then thaw-mounted onto ITO-coated electrically conductive glass slides. The serial sections from the tissues that would be compared should be thraw-mounted onto the same ITO slide, acquired and normalized together.

The method was developed based on our previously described method. In briefly, α-cyano-4-hydroxycinnamic acid (CHCA, No. C2020, Sigma-Aldrich) was used as a MALDI matrix in positive ion modes, 9-aminoacridine (9AA, No. 92817, Sigma-Aldrich) was used as a MALDI matrix in negative ion modes. A two-step method was performed for the matrix applications by combining sublimation and airbrushing(*82*). The first step is sublimation. The ITO-coated glass slide bearing the brain tissue specimen was installed in a sample holder and was embedded in a vacuum deposition system (SVC-700TMSG iMLayer, Sanyu Electron, Tokyo, Japan). A matrix holder filled with approximately 300 mg of matrix powder was positioned 8 cm from the sample holder. The matrix powder was then heated to the boiling point of the matrix crystals (9AA, 220 °C; CHCA, 250 °C). The vapor was allowed to cover the specimen surface. The vacuum pressure of the chamber was maintained at 10-4 Pascal (Pa) during the sublimation procedure. The second step was airbrushing. Briefly, matrix solution (10 mg/mL of 9AA or CHCA) was prepared by dissolving matrix powder in acetonitrile and distilled water containing 0.1% FA at a ratio of 1:1 (v/v). One milliliter of the matrix solution was added to the capacity of an artist’s airbrush (MR Linear Compressor L7/PS270 Airbrush, GSI Creos, Tokyo, Japan) and sprayed onto the surface of tissues for 23 cycles. The distance between the tip of the airbrush and the tissue surface was approximately 8 cm. During the period of the first 3 cycles, the matrix was airbrushed for 2 s at 60 s intervals. After that, the matrix was continuously sprayed for 1 s at 30 s intervals during the following 20 cycles. The sample slide was then placed in a vacuum desiccator for 5 min to vaporize the solvent.

MALDI IMS was performed on a well-established record of high resolution and precise mass measurement capabilities due to its IT-TOF detector combining ion trap and TOF. High-resolution, high-precision fine mass measurement can be done in MS, MS/MS, MS/MS/MS mode due to the Dual-stage Reflectoron (DSR) and RF Ballistic Ion Extraction (BIE) technology. The ions were identified using the mass-charge ratio (m/z) as the x-axis and the ion abundance as the y-axis. The MALDI source uses a diode-pumped 355 nm Nd:YAG laser (Shimadzu Corporation, Kyoto, Japan). The relevant region of the tissue was chosen using an optical microscope embedded in the iMScope prior to data acquisition. A tissue ROI was selected via a charge-coupled device (CCD) camera (Olympus Corporation, Tokyo, Japan). The illumination in the iMScope was applied according to the following parameters: light type, transillumination; light intensity, 12%. The MS acquisition parameters were set as follows: ion polarity, positive (after coated with CHCA) or negative (after coated with 9AA); mass range, 200–400 (ion polarity, positive); 200-700 (ion polarity, negative); sample voltage, 3.5 kV (ion polarity, positive); 3 kV (ion polarity, negative); detector voltage, 1.90 kV. Laser firing parameters: frequency, 1000 Hz; laser intensity, 55 arbitrary unit of iMScope; laser diameter, 3 arbitrary unit of iMScope. All the experiments in this work were performed with the minimum irradiation diameter to irradiate the tissue surface with 100 shots for each pixel. The imaging MS Solution Version 1.30 software (Shimadzu, Tokyo, JPN) was used to operate the instrument, and the data acquisition, visualization, and quantification were also performed using the same software. According to the mass-to-charge ratio (m/z) obtained from MALDI-MSI, individual small molecules were identified in the HMDB database (https://hmdb.ca) or Lipid Maps database (https://www.lipidmaps.org/) and compared with commercially available small molecules standards. Additional confirmation of small molecules identities was performed according to comparisons with published reports and based on the ion spectra of authentic standards by MALDI-TOF-MS/MS assays. The m/z values were internally calibrated with 2,5-dihydroxybenzoic acid (DHB No. 149357, Sigma-Aldrich).

### Immunofluorescence (IF) and Immunohistochemistry (IHC)

For immunofluorescence, tissue samples were fixed in 4% paraformaldehyde, cryoprotected in 30% sucrose, embedded in Optimal Cutting Temperature (OCT) compound (Sakura Finetek), and sectioned at 10 µm thickness using a cryostat. Sections were permeabilized with 0.1% Triton X-100 in PBS for 15 minutes and blocked with 5% normal donkey serum containing 1% bovine serum albumin in PBS for 1 hour at room temperature. The sections were then incubated with primary antibodies diluted in blocking solution overnight at 4°C. After washing with PBS containing 0.05% Tween-20 (PBST), the sections were incubated with appropriate species-specific secondary antibodies conjugated with Alexa Fluor 488 or 594 (Abcam, 1:400) for 1 hour at room temperature in the dark. Nuclei were stained with 4’,6-diamidino-2-phenylindole (DAPI; Solarbio, cat# C0060) for 5 minutes. Slides were mounted with anti-fade mounting medium (Servicebio, cat# G1401) and imaged using a confocal laser scanning microscope (LSM 900, Zeiss). Images were processed and analyzed using ZEN software.

For IHC, tissue samples were fixed in 4% paraformaldehyde, embedded in paraffin, and sectioned at 5 µm thickness. After deparaffinization and rehydration, heat-induced epitope retrieval was performed in sodium citrate buffer (10 mM, pH 6.0) using a microwave. Subsequent immunohistochemical staining was performed using a two-step IHC detection kit (Maxim Biotechnologies; KIT-9701 for rabbit primary antibodies, KIT-9706 for mouse primary antibodies) according to the manufacturer’s instructions. Briefly, endogenous peroxidase activity was quenched with 3% hydrogen peroxide for 15 minutes at room temperature. Sections were blocked with the provided blocking reagent for 30 minutes and then incubated with primary antibodies overnight at 4°C in a humidified chamber. After washing, the sections were incubated with the kit’s ready-to-use biotinylated secondary antibody for 30 minutes at 37°C, followed by horseradish peroxidase-conjugated streptavidin for 30 minutes. The immunoreactivity was visualized using 3,3’-diaminobenzidine (DAB) chromogen (Maxim, DAB-0031). Nuclei were counterstained with hematoxylin for 30 seconds. Finally, slides were dehydrated through graded ethanol series, cleared in xylene, and mounted with neutral balsam. Images were acquired using a bright-field microscope (ECLIPSE Ci-L, Nikon) and analyzed with ImageJ software.

Primary antibodies used for IHC:

Rabbit anti-IBA1 (Wako, cat# 019-19741, 1:400)

Rabbit anti-NeuN (Abcam, cat# ab177487, 1:400)

Rabbit anti-PD-L1 (Abcam, cat# ab205921, 1:200)

Rabbit anti-CD39 (Abcam, cat# ab227840, 1:200)

### Western Blot Analysis

Protein lysates were extracted from primary bone marrow macrophages and tissues using RIPA lysis buffer (Beyotime, P0012B) supplemented with protease (MedChemExpress, HY-K0010) and phosphatase inhibitors (MedChemExpress, HY-K0021). Protein concentrations were determined using a bicinchoninic acid (BCA) assay. Equal amounts of protein (typically 20–40 µg per lane) were separated by SDS-polyacrylamide gel electrophoresis (SDS-PAGE) and subsequently transferred onto polyvinylidene fluoride (PVDF) membranes (Merck Millipore, IPVH00010). After blocking, membranes were incubated overnight at 4 °C with specific primary antibodies. Following incubation with appropriate horseradish peroxidase (HRP)-conjugated secondary antibodies for 1 h at room temperature, immunoreactive bands were visualized using an enhanced chemiluminescence (ECL) substrate and captured with a chemiluminescence imaging system (Tanon, 5200). Band intensities were quantified using ImageJ software.

Primary antibodies used in this study:

Mouse anti-CDKN2A/p16 (Santa Cruz Biotechnology, cat# sc-1661, 1:1000)

Mouse anti-CDKN1A p21 (Santa Cruz Biotechnology, cat# sc-6246, 1:1000)

Mouse anti-p53 (Santa Cruz Biotechnology, cat# sc-98, 1:1000)

Mouse anti-GAPDH (Absin, abs830030,1:10000)

Rabbit anti-γH2A.X (Cell Signaling Technology, cat# 9718S, 1:1000)

### RNA extraction and RT-qPCR

Total RNA was extracted from cells or tissue using SteadyPure Quick RNA Extraction Kit (Cat#: AG21023, Accurate Biology) according to the manufacturer’s instructions. cDNA templates were synthesized using PrimeScript™ RT reagent Kit with gDNA Eraser for qPCR (+gDNA wiper) (Cat#: RR047A, Takara). RT-qPCR assays were performed with TB Green® Premix Ex Taq™ II (Cat#: RR820A, Takara) and LightCycler® 96 Instrument. Relative mRNA level of the indicated gene was normalized to ACTB. The following primers were used: *Gzmk* (F): 5’ ACCGTGGTTTTAGGAGCACAT-3’; *Gzmk* (R): 5’-CCCAGGTGAAGCAGTTGGAC; *Atcb* (F): GGCTGTATTCCCCTCCATCG-3’, *Actb* (R): 5’-CCAGTTGGTAACAATGCCATGT- 3’; *Cdkn2a* (F) 5′-CCCAACGCCCCGAACT*-3*′*, Cdkn2a* (R) 5′- GCAGAAGAGCTGCTACGTGAA-3′; Firefly luciferase (F) 5′- GCCATGAAGCGCTACGCCCTGG-3′, luciferase (R) 5′-TCTTGCTCACGAATACGACGGTGG-3′.

### Quantification and statistical analysis

Statistical analyses were conducted using Prism software 9.0 (GraphPad). Data are expressed as means with error bars representing the standard error of the mean (SEM), unless otherwise indicated. For two-group comparisons, paired or unpaired two-tailed Student’s t-tests were used.

For multiple group comparisons, either two-way ANOVA with post-hoc Tukey, Šídák, or Dunnett tests, or one-way ANOVA with post-hoc Tukey or Dunnett tests, were applied as appropriate. A p-value of less than 0.05 was considered statistically significant. Significance levels are denoted as ∗p < 0.05, ∗∗p ≤ 0.01, ∗∗∗p ≤ 0.001, and ∗∗∗∗p ≤ 0.0001, and specific details are provided in the figure legends.

## Acknowledgments

This work was supported by the National Key Research and Development Program of China 2023YFC3606600 (to W.C.), the National Natural Science Foundation of China 82571797, 32270938 (to W.C.), the Public Hospital Research Collaboration Fund of Inner Mongolia (2023GLLH0336 to K.X.).

## Author contributions

W.C., ZY.W., Z.W. and Y.L. designed the study. L.G., R.Z., Q.Z. and X.Y. performed experiments. Q.Z. performed IMS experiment. J.Y. analyzed the scRNA-seq. Z.W., ZY.W., Y.L. and W.C.analyzed and interpreted data. ZY.W. and W.C. wrote the manuscript with input from all authors.

