## Supplemental figures for "A feed-forward loop between niche adenosine and Gzmk⁺ CD8 T cells propagates systemic inflammaging"

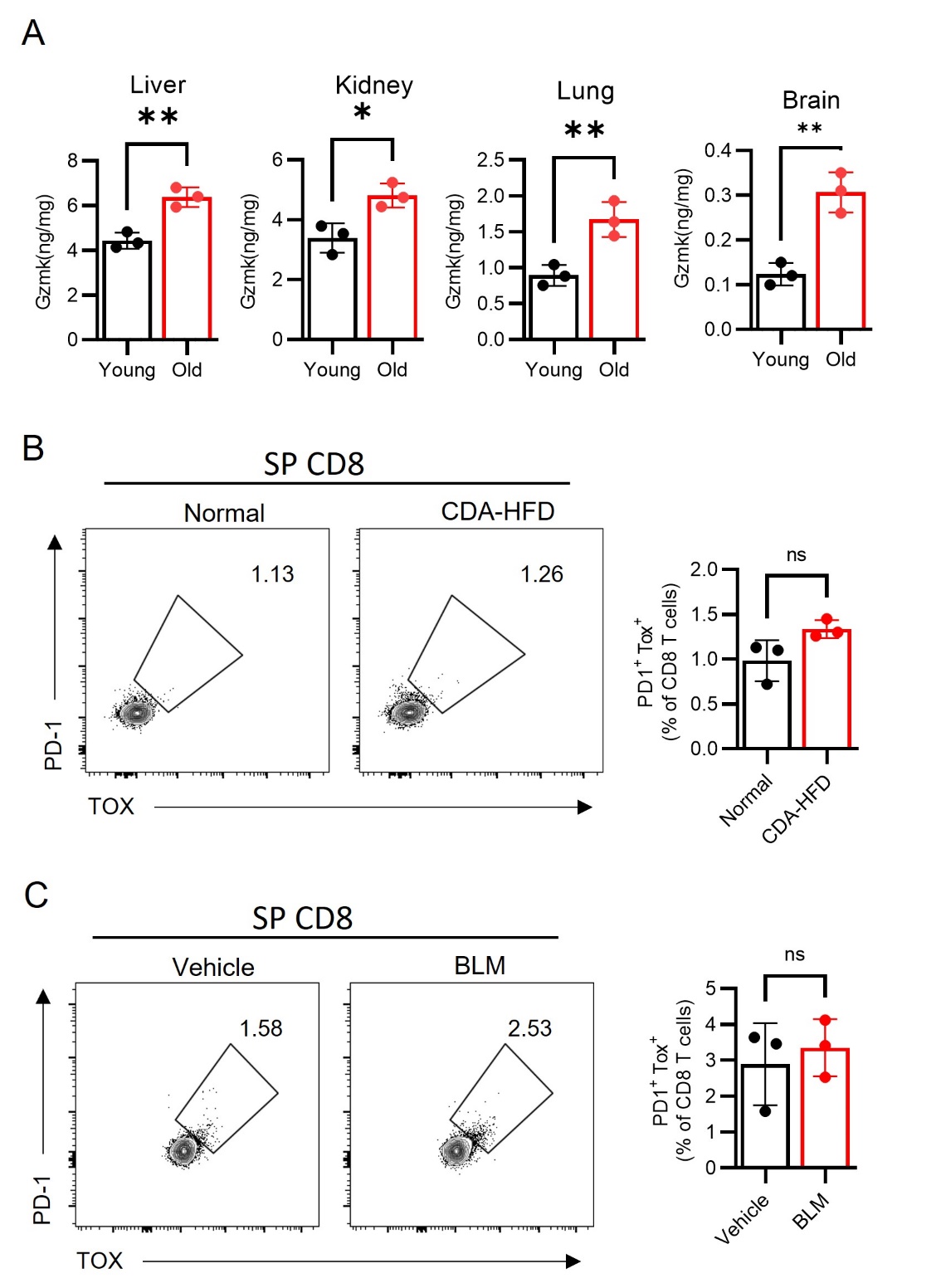

**Figure S1, Related to Figure 1, Gzmk^+^ CD8⁺ T cells are generated locally in response to organ-specific aging.** (A) Gzmk level in indicated tissues were measured by ELISA. CD8 T cells from spleen were analyzed by flow cytometry in Figure 1G (B), Figure 1H (D).

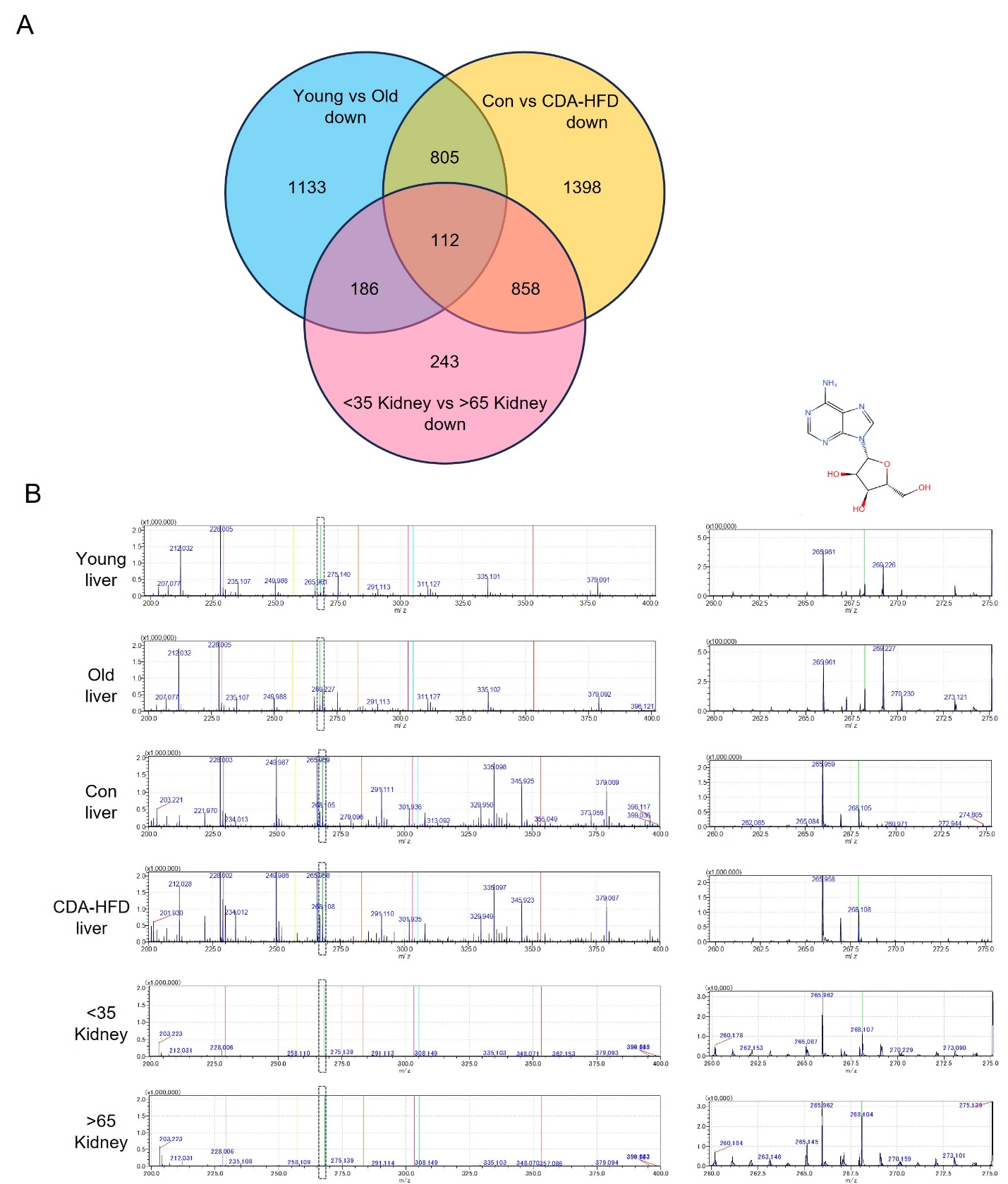

**Figure S2, IMS analysis of indicated tissues from young and old mice or young and old individuals or control and CDA-HFD-feed mice.** (A) Identification of metabolites upregulated at livers from old and CDA-HFD-feed mice and kidneys from old individuals. (B) Signaling peaks from m/z 200 - m/z 400 at indicated tissues.

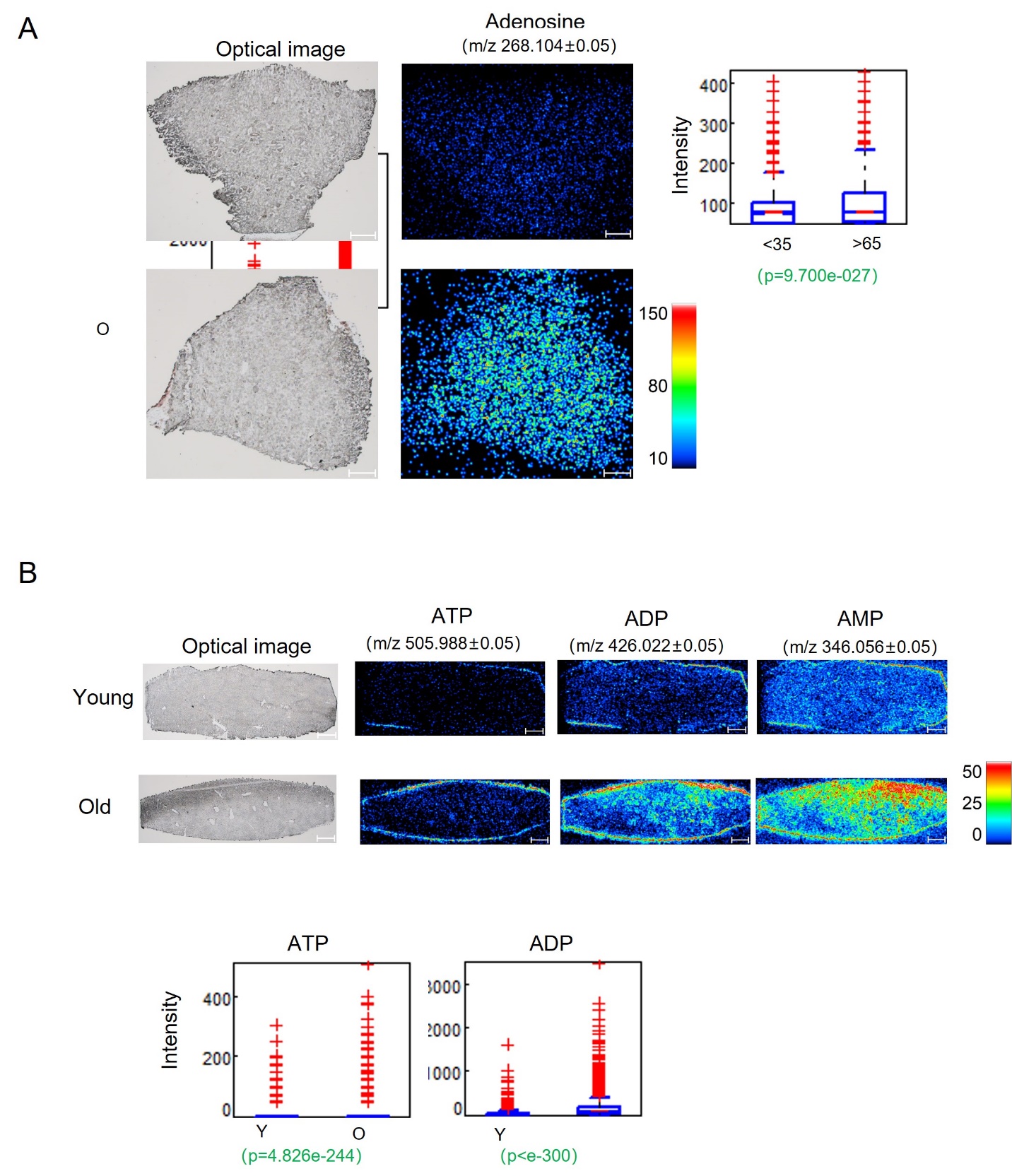

**Figure S3, IMS analysis of young and old tissues.** (A) Visualization of adenosine signaling of kidney from young and old individuals revealed by IMS analysis. (B) Visualization of ATP, ADP and AMP signaling of liver from young and old individuals revealed by IMS analysis.

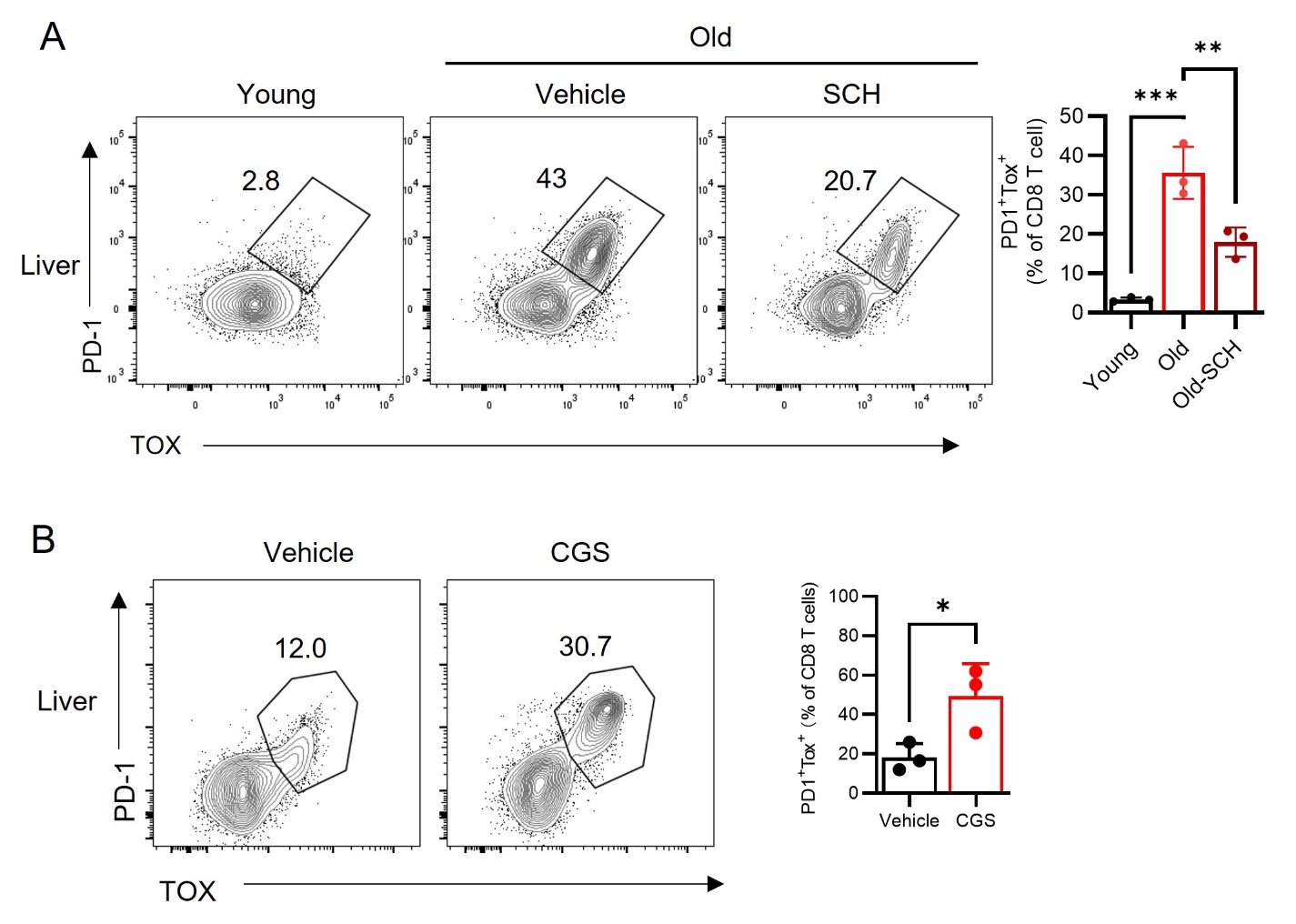

**Figure S4, related to Figure2. Adenosine accumulation in aged tissues drives Gzmk⁺ CD8 T cell differentiation via A2aR–Eomes signaling*.*** PD1^+^ TOX^+^ CD8 T cells in liver were analyzed by flow cytometry in Figure 2D (A). PD1^+^ TOX^+^ CD8 T cells in liver and brain were analyzed by flow cytometry in Figure 2C (B).

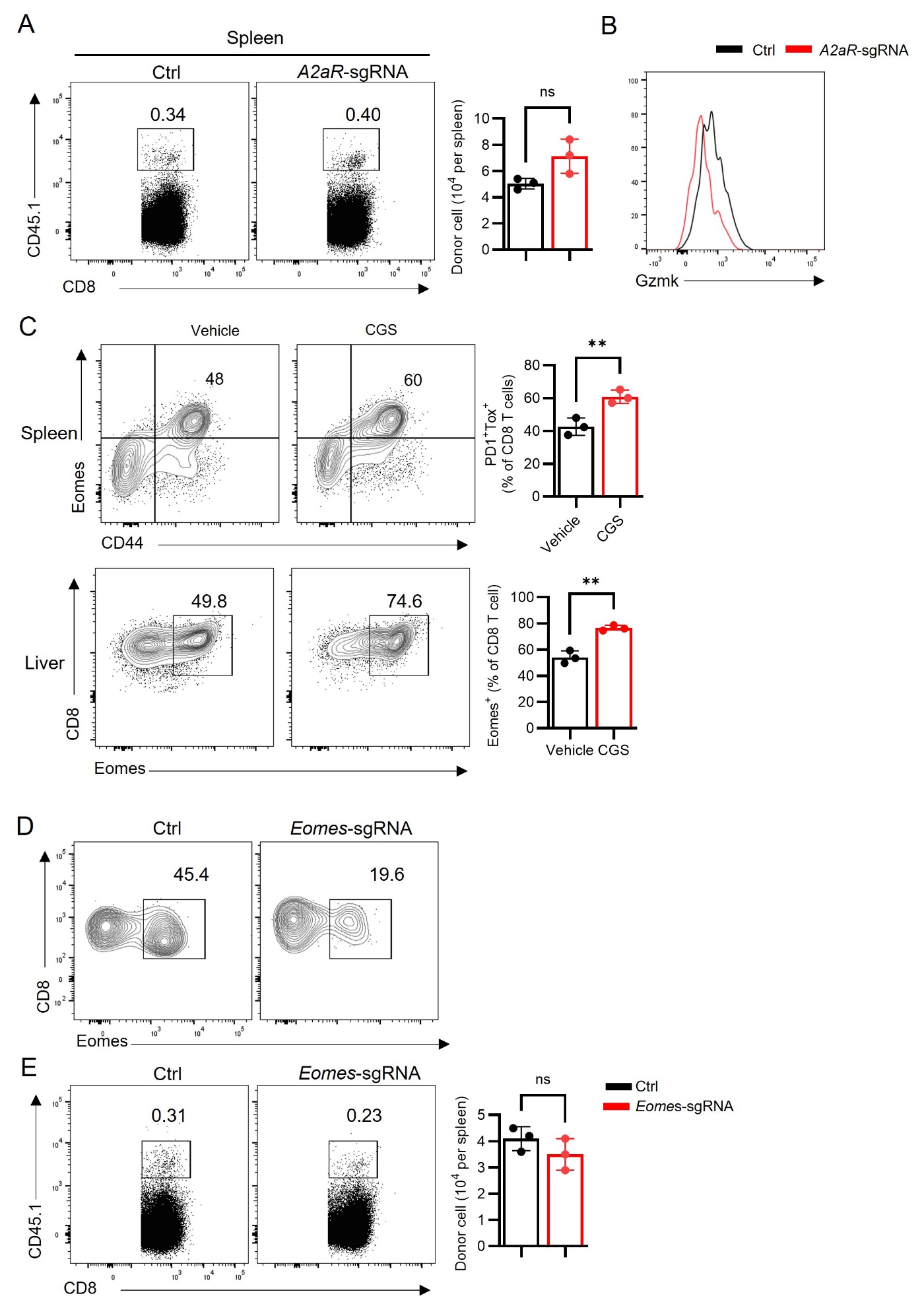

**Figure S5, related to Figure2. Adenosine accumulation in aged tissues drives Gzmk⁺ CD8 T cell differentiation via A2aR–Eomes signaling*.*** The percentage and cell number of donor cells (A), the expression of Gzmk in donor cells analyzed by flow cytometry (B) in Figure 2F. (C) Eomes expression in CD8 T cells in spleen and liver from young or old mice were analyzed by flow cytometry in Figure 2E. Efficiency of *Eomes* knock-out analyzed by flow cytometry (D), the percentage and cell number of donor cells (E) in Figure2H.

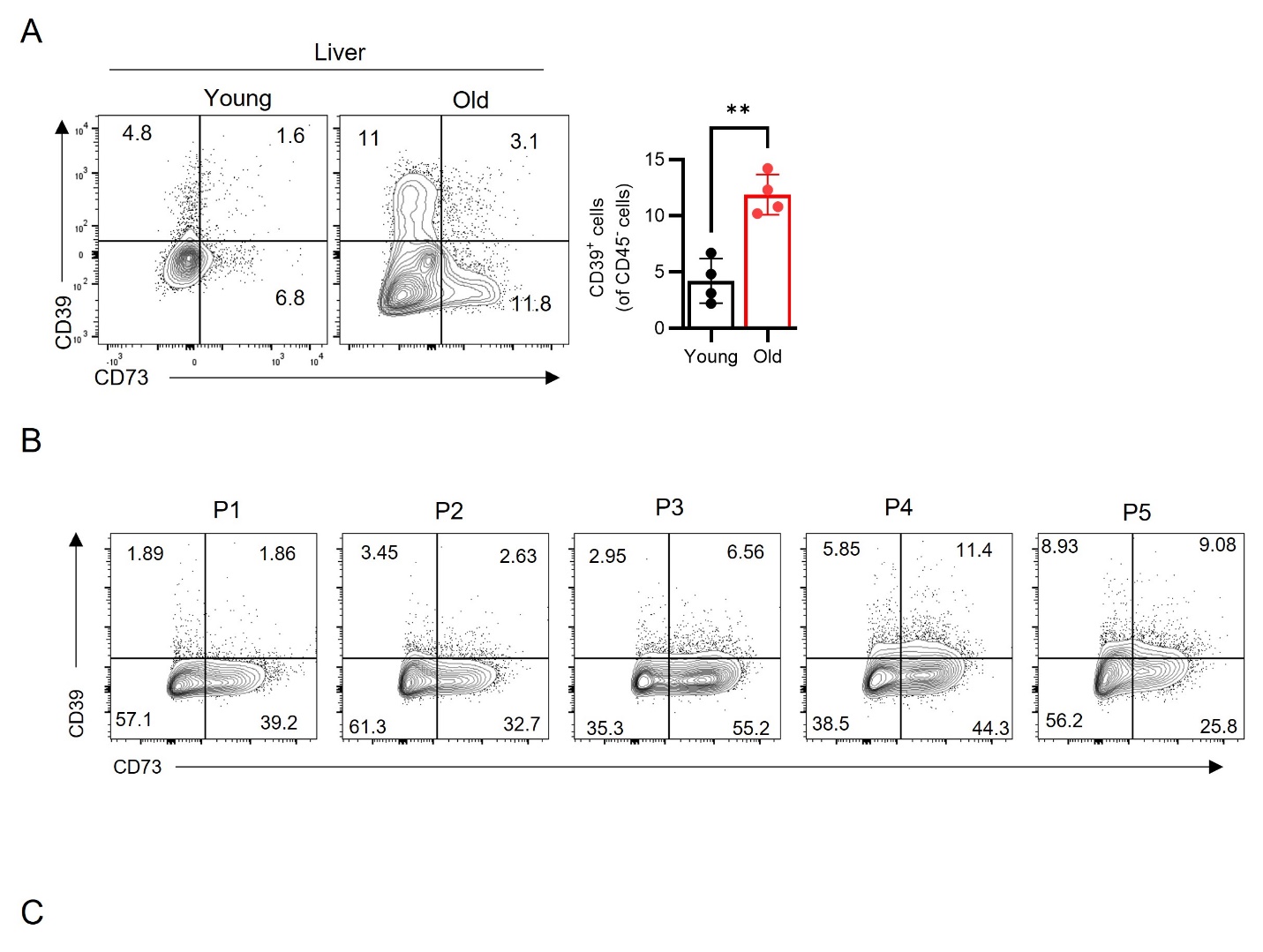

**Figure S6, related to Figure 3, p16-dependent CD39 induction in senescent cells drives adenosine accumulation.** (A) Expression of CD39 and CD73 on CD45^-^ cells in liver from young or old mice were analyzed by flow cytometry. (B) MEFs were continuously passaged and CD39 and CD73 were determined by flow cytometry. (C) The targeting strategy of *p16^Ink4a^*-luciferase reporter mice

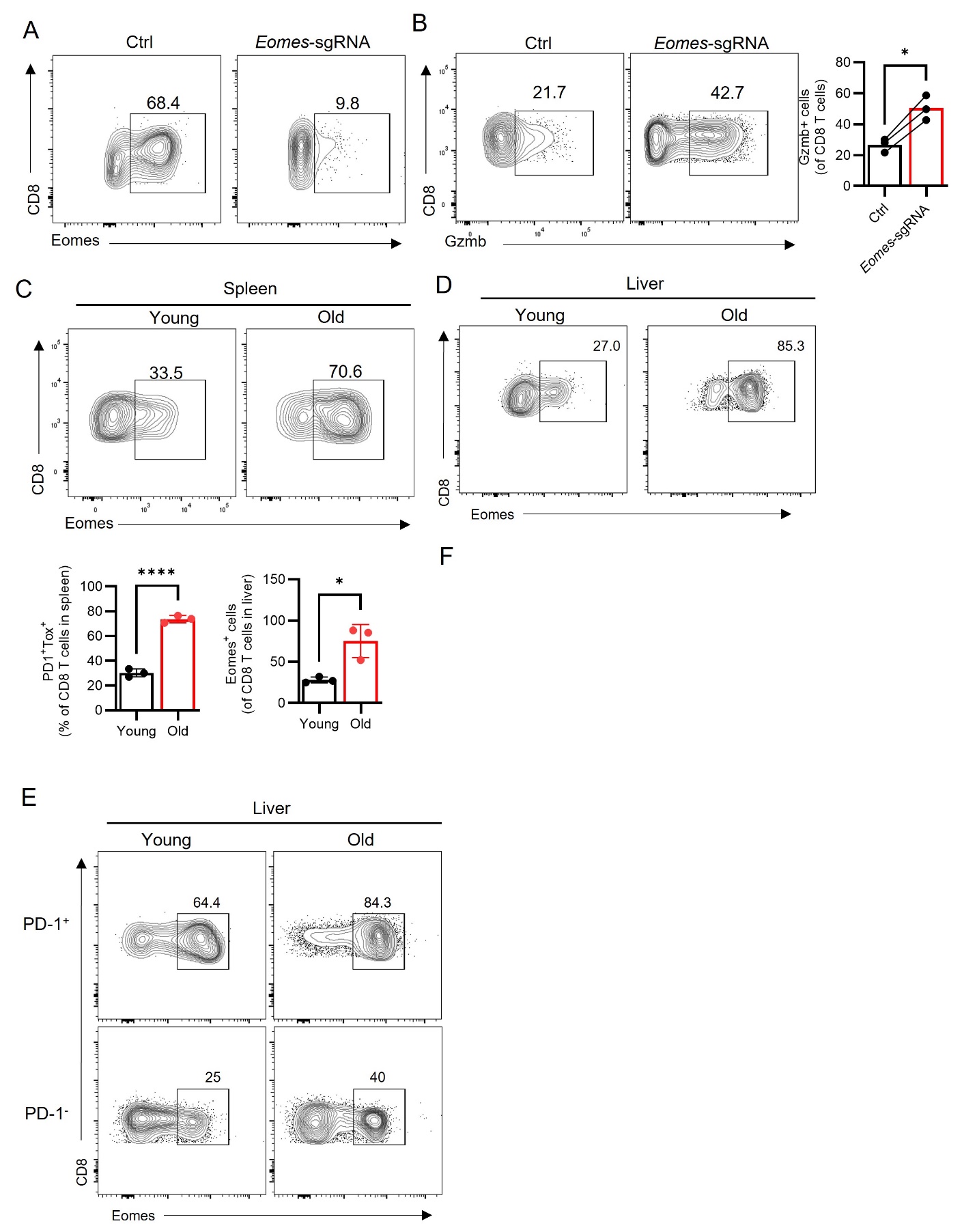

**Figure S7, related to Figure 4, Eomes regulates Gzmk in response to adenosine and PD-L1 stimulation.** (A-B) Related to Figure 4B, Cas9-mediated Eomes knock-out was confirmed by flow cytometry (A), production of Gzmb was determined by flow cytometry (B). Eomes expression of CD8 T cells in spleen (C), in liver (D) from young or old mice were analyzed by flow cytometry. (E) Representative contour plots of CD8 T cells in Figure 4C.

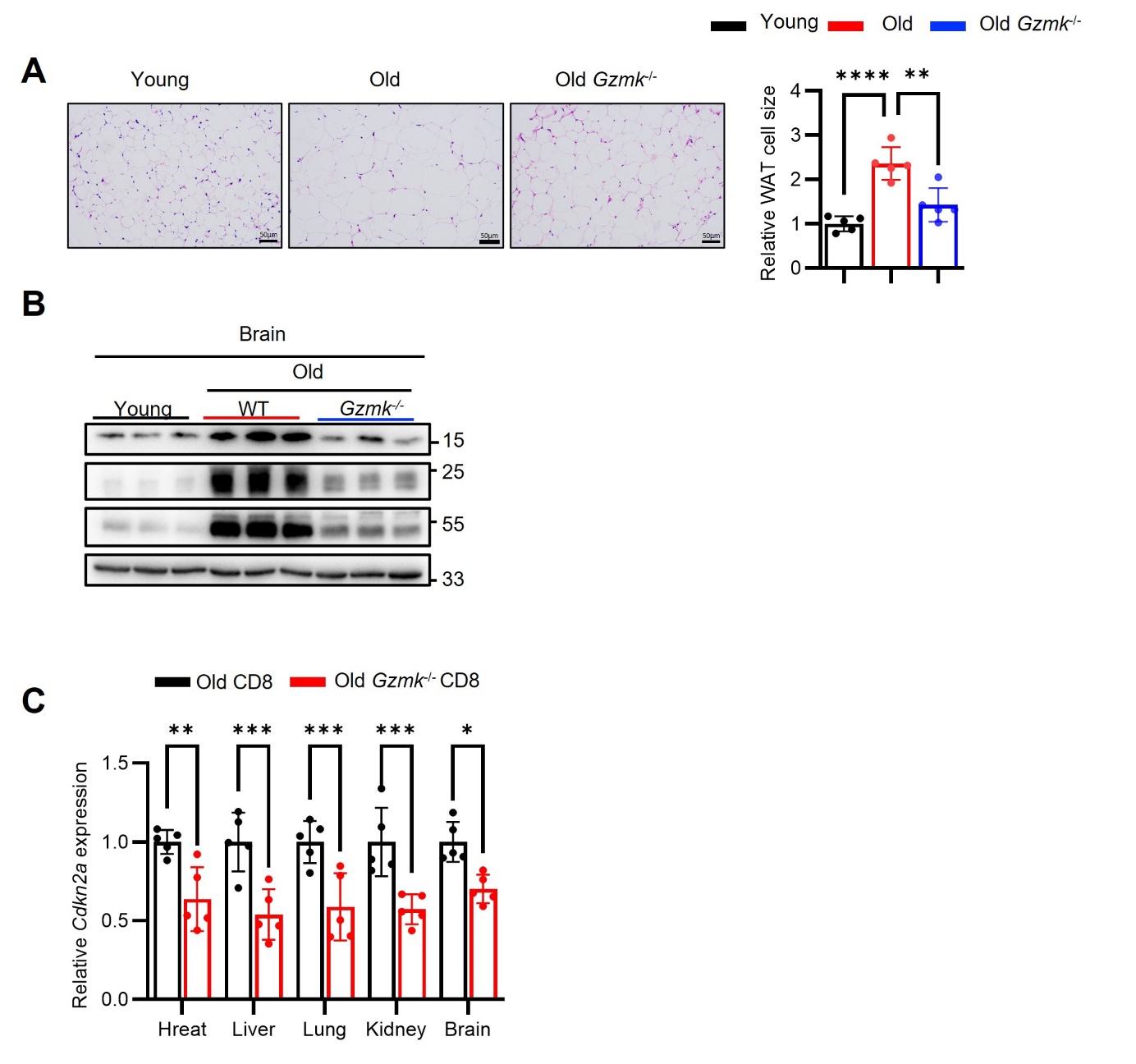

**Figure S8, related to Figure 6. CD8 T cell-derived Gzmk triggers systemic cellular senescent across organs.** (A, B) related to Figure 6A-D. (A) H&E staining for WAT from indicated mice. (B) Western blot analysis of P16, P21 and P53 in brain. (C) related to Figure 6F-G, the expression of *Cdkn2a* in indicated tissues were measured by RT-qPCR.

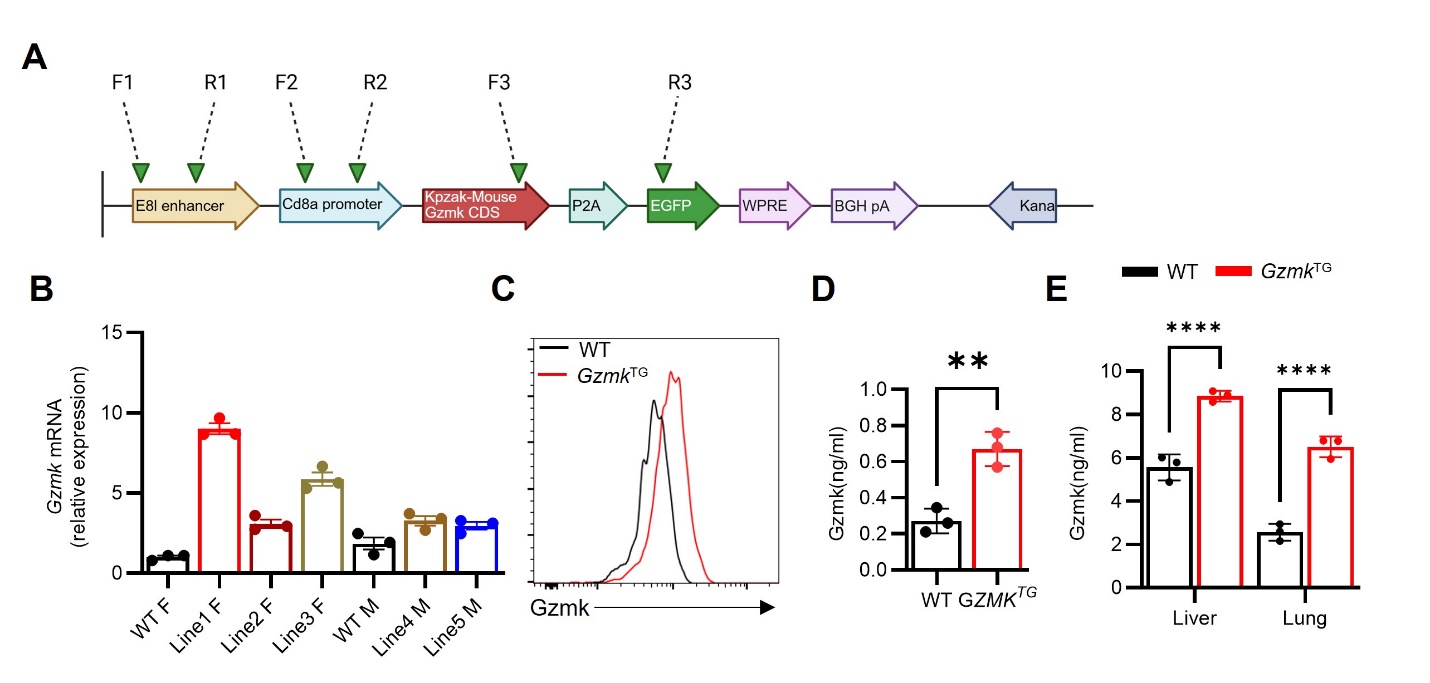

**Figure S9. Generation and screen of *Gzmk*^TG^ mice** (A) The generation strategy of Gzmk transgenic mice. (B) qPCR analysis of splenocytes from different founder lines. (C-F) Verification of line 1 (referred to as *Gzmk*^TG^), Gzmk expression in purified CD8 T cells from spleen (C), level of Gzmk in plasma measured by ELISA (D), level of Gzmk in indicated tissues measured by ELISA

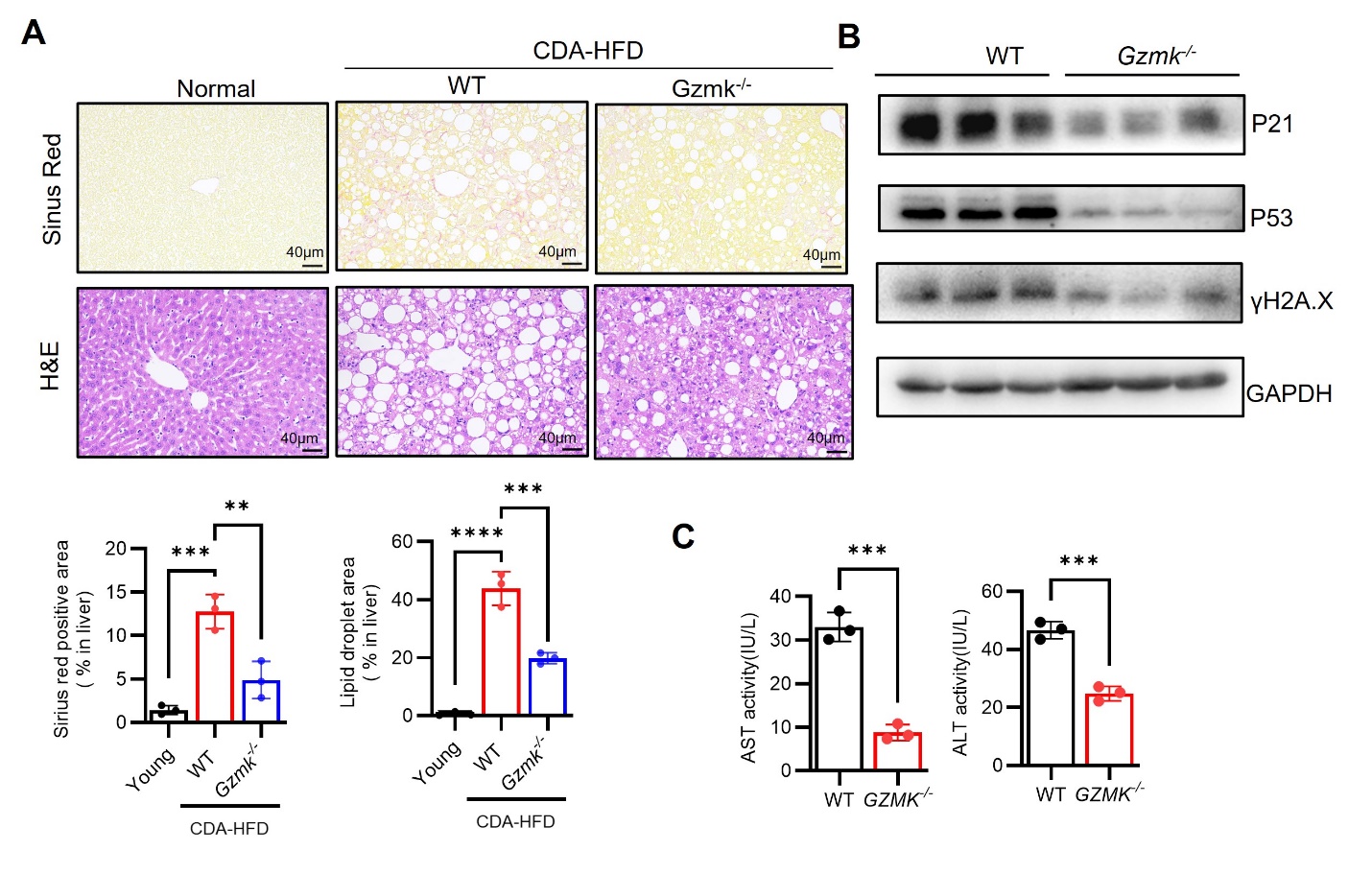

**Figure S10. Genetic ablation of *Gzmk* confers protection against senescence-associated NASH pathology.**  WT or *Gzmk^-/-^* mice were subjected to NASH induction. (A) Staining of Sinus Red and H&E of livers from indicated mice. (B) Western blot analysis of the expression of p16, p21 and p53 in liver. (C) AST and ALT in plasma from indicated mice were measured by ELISA.

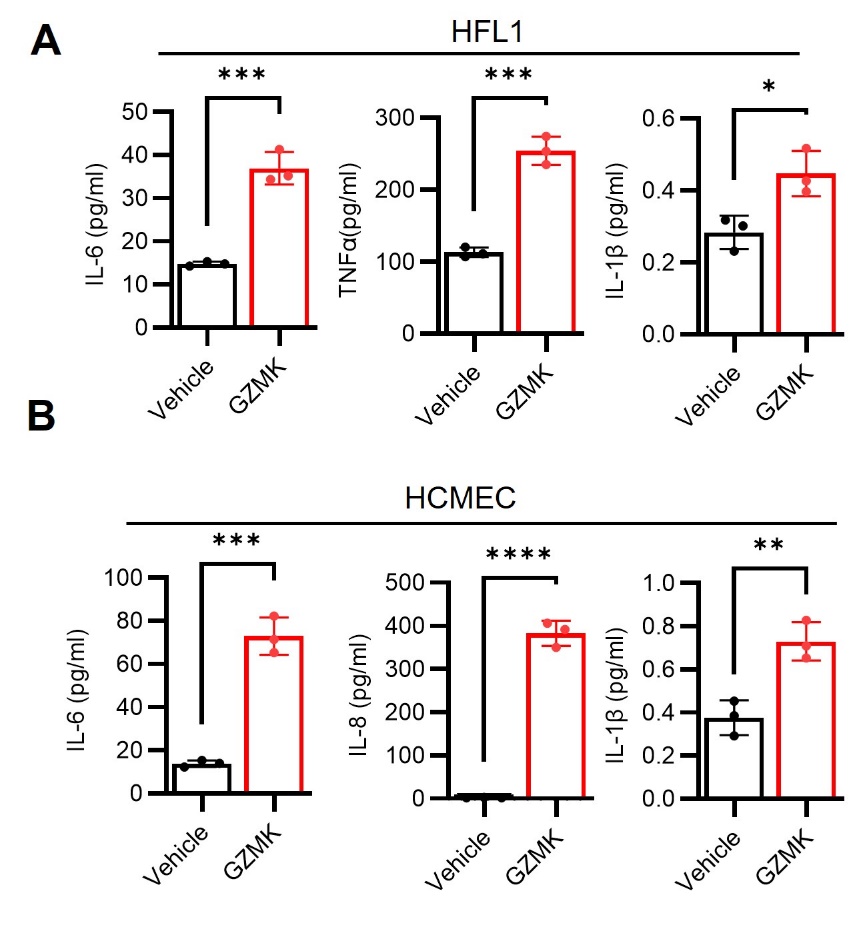

**Figure S11, related to Figure 8. Extracellular Gzmk Drives Cellular Senescence and Inflammaging via a PAR1-MAPK Axis.** HFL1 (A), HCMEC (B) were treated with 100nM GZMK in medium without FBS for one day, IL-6, IL-8, IL-1β and TNFα in supernatant were measured by ELISA.

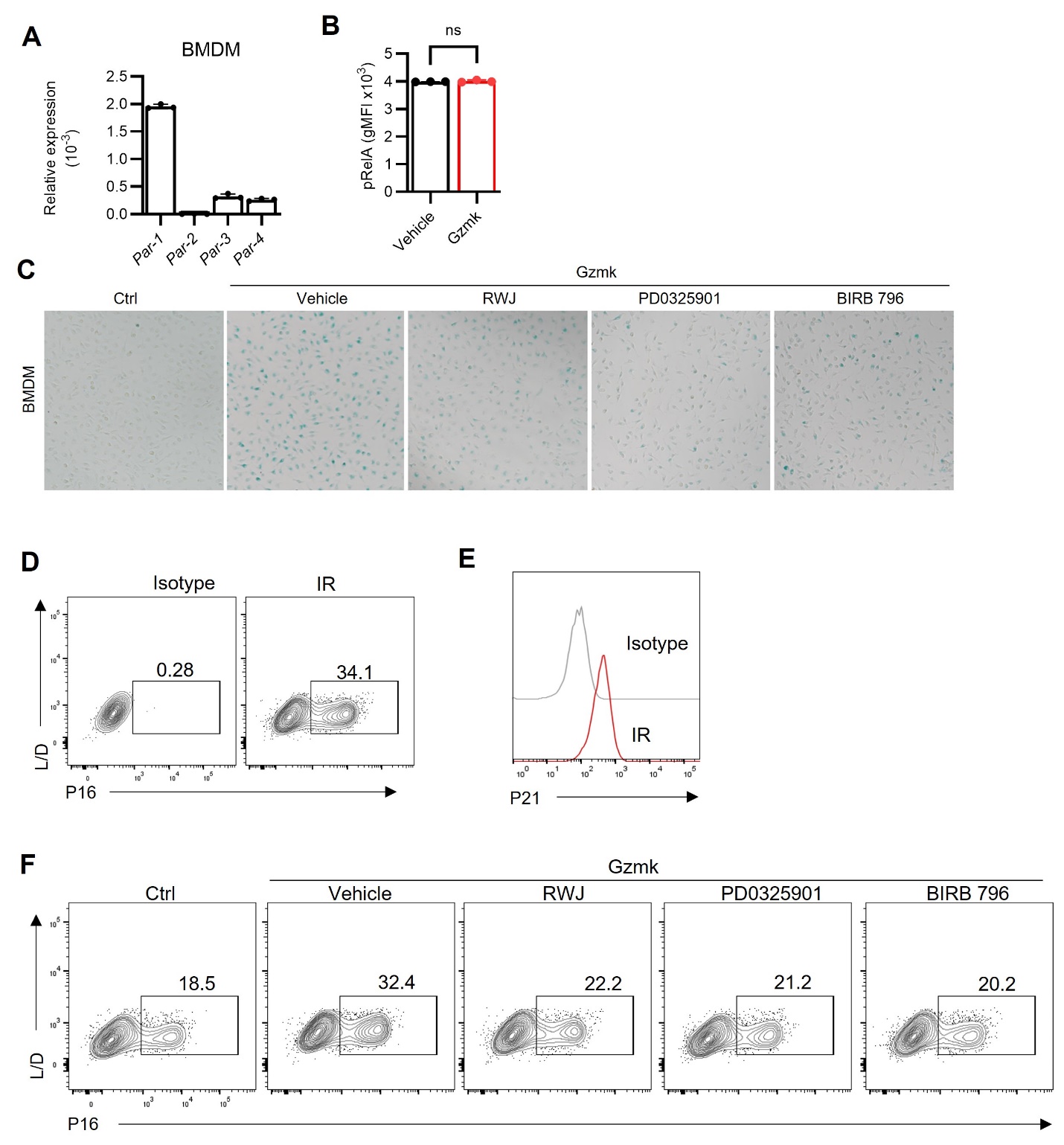

**Figure S14, related to Figure 8. Extracellular Gzmk Drives Cellular Senescence and Inflammaging.** (A) qPCR analysis for *Par-1/2/3/4* expression in BMDM. (B) Phosphorylated RelA in Figure 8C was determined by flow cytometry. (C) The representative SA-β-gal staining in Figure 8D. (D) The validation of p16 staining in BMDM by flow cytometry. (E) The representative dot contour flow data in Figure 8F. (F) The validation of p21 staining in BMDM by cytometry.

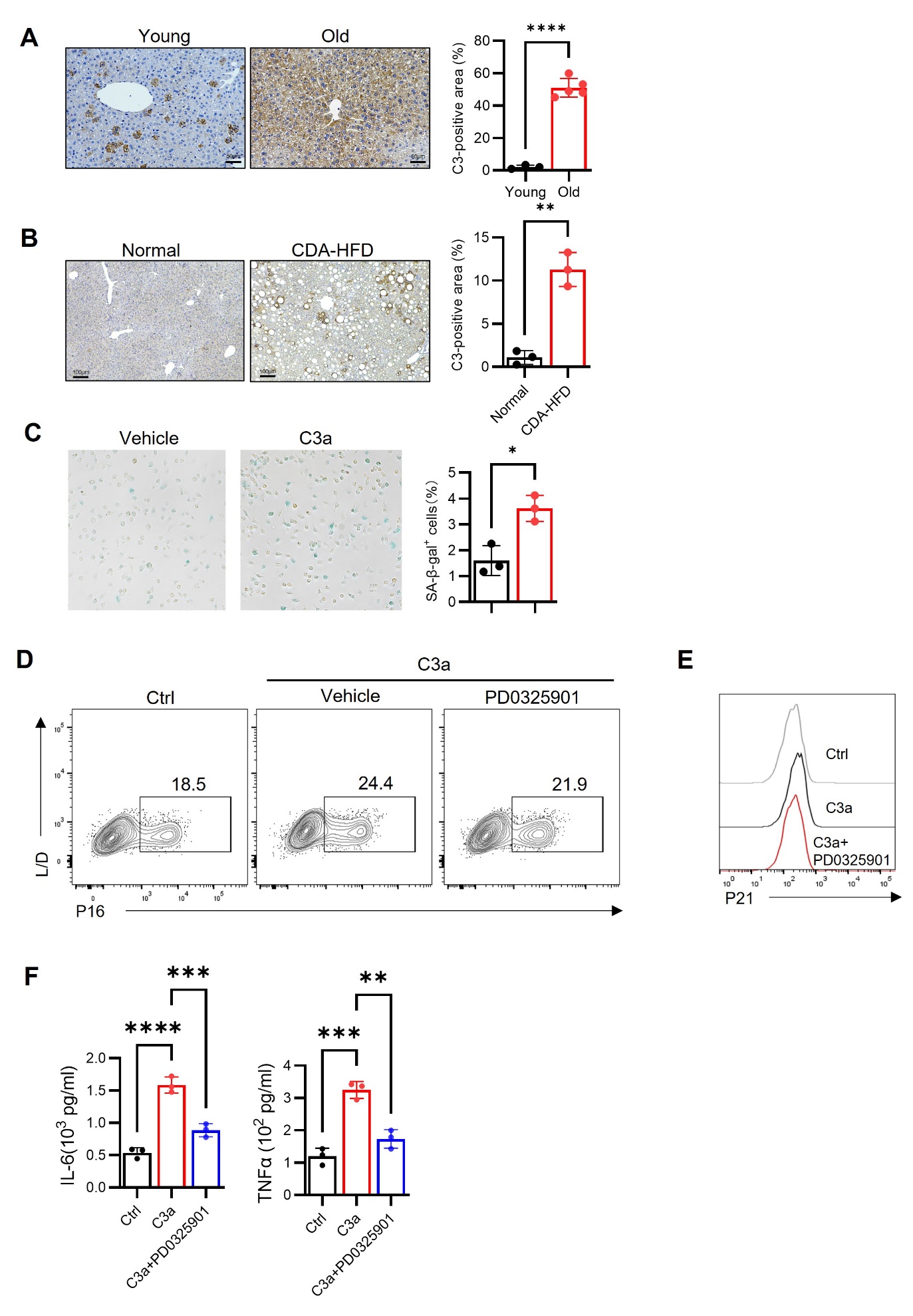

**Figure S13. C3a Contributes to Cellular Senescence and Inflammaging.**  C3 staining in liver from young or old mice (A), control or NASH mice (B). (C) BMDM were treated with C3a for three days, and SA-β-gal staining was performed. (D, E) BMDM were stimulated with C3a for one day, then P16 and P21 expression were measured by flow cytometry. BMDM were treated with C3a or combined with MEK inhibitor in medium without FBS for one day, IL-6 TNFα in supernatant were determined by ELISA.
